# Shh induces symmetry breaking in the presomitic mesoderm by inducing tissue shear and orientated cell rearrangements

**DOI:** 10.1101/589549

**Authors:** J. Yin, T. E. Saunders

## Abstract

Future boundaries of skeletal muscle segments are determined in the presomitic mesoderm (PSM). Within the PSM, future somitic cells undergo significant changes in both morphology and position. How such large-scale cellular changes are coordinated and the effect on the future border formation is unknown. We find that cellular rearrangements differ between cell populations within the PSM. In contrast to lateral somitic cells, which display less organized rearrangement, the adaxial cell layer undergoes significant tissue shearing with dorsal and ventral cells sliding posteriorly. This shear is generated by orientated intercalations of dorsally and ventrally located adaxial cells, which induces a chevron-like pattern. We find Shh signaling is required for the tissue shear and morphogenesis of adaxial cells. In particular, we observe Shh-dependent polarized recruitment of non-muscle myosin IIA drives apical constrictions, and thus the intercalations and shear. This reveals a novel role for Shh in regulating cell mechanics in the PSM.

## Introduction

Spatial patterns during development are often determined through signaling molecules called morphogens (Gurdon and Bourillot, 2001; Turing, 1952). A classic example is patterning of the anterior-posterior (AP) axis in the early *Drosophila* embryo by Bicoid, Caudal, and Nanos (Driever et al., 1989; Frohnhöfer and Nüsslein-Volhard, 1986; Irish et al., 1989). Considerable work has identified how the positional information from morphogens is interpreted to ensure robust gene expression boundaries (Dubuis et al., 2013; Gregor et al., 2007; Lander, 2013). However, in many tissues, the system is undergoing morphological changes during patterning (Kicheva et al., 2014; Xiong et al., 2013). It remains unclear how signaling pathways and mechanical interactions interplay to ensure robust formation and patterning of tissues. To explore this general problem, here we focus on the formation of vertebrate future muscle segments.

The metameric structures of the vertebrate body, including the axial skeleton and skeletal muscles, are derived from mesodermal building blocks within embryos called somites (Dequéant and Pourquié, 2008; Hubaud and Pourquié, 2014). The somites, positioned on the lateral sides of the embryonic midline, are periodically segmented from the presomitic mesoderm (PSM) through a “clock and wavefront” mechanism (Baker et al., 2006; Cooke and Zeeman, 1976; Krol et al., 2011). In this model, the clock (a series of molecular oscillators) and the wavefront (a posteriorly moving signaling gradient that arrests the clock) determine the timing and position of the emerging somite border. The earliest boundaries of future somites are specified within the PSM by the fibroblast growth factor (FGF) (Dubrulle et al., 2001; Sawada et al., 2001). This specification occurs around S-5 in the tailbud of the zebrafish embryo, *i.e.* around 200 minutes before that somite is segmented from the PSM (given 40 minutes period at 26°C used in this study) (Akiyama et al., 2014). Thus, spatial patterning of boundaries within the PSM occurs before the somite boundaries themselves are clearly delineated. The polarized expression of cMeso-1 in posterior border cells of chick embryos (vertebrate Mesp2) upregulates Eph/Ephrin signaling, leading to deposition of extracellular matrix proteins between the inter-somitic boundaries (Barrios et al., 2003; Julich et al., 2009; McMillen et al., 2016; Watanabe et al., 2009).

Cell sorting is an active mechanism of cell rearrangements, dependent on the cell mechanical properties. Cell sorting has been observed in the zebrafish embryo neural tube, where mixed populations of progenitor cells are separated and subsequently their domain boundaries sharpen (Xiong et al., 2013). It is not clear whether similar sorting is occurring in the PSM to regulate future somite boundaries. Recently, a fluid-to-solid jamming transition was observed across the tailbud and PSM (Mongera et al., 2018). Cells migrate more rapidly in the tailbud and posterior PSM, whilst cells located in the anterior PSM are more rigidified. Given these large-scale morphological changes, it remains an open question as to how the future somite boundaries – determined 5 cycles before somite segmentation - are robustly maintained prior to segmentation.

There are two major cell populations within the PSM, determined according to their locations, morphologies and future cell fates: the medial cells and lateral cells (Devoto et al., 1996). The medial cells, a planar array of cells with epithelial morphology, are located at the medial side of somite, and are mainly composed of adaxial cells located along the midline in close contact to the notochord (Hirsinger et al., 2004). In the somite, the adaxial cells differentiate into slow twitch muscles, a process dependent on Sonic Hedgehog (Shh) signalling from the notochord. The final cell fate of the more dorsally and ventrally located medial cells remains unclear (Daggett et al., 2007; Yin et al., 2018). The lateral cells of the PSM, the majority of which display a mesenchymal morphology, differentiate into fast twitch muscles, dermomyotome and non-muscle progenitors after somite segmentation (Hollway et al., 2007; Stellabotte et al., 2007; Yin et al., 2018). Shh, Bone Morphogenetic Protein (BMP), and Fibroblast Growth Factor (FGF) signalling determines cell fate within both adaxial and lateral cell lineages (Maurya et al., 2011; Nguyen-Chi et al., 2012; Wolff et al., 2003; Yin et al., 2018). However, it is unknown how these signalling pathways interplay with cell morphological changes during somite formation to ensure robust boundary formation. In particular, how, if at all, do these signalling pathways regulate cell mechanical properties during somite formation?

In this study, by utilizing cell tracking from extended live imaging of the PSM and somites, we identified the future somite boundaries within the PSM as early as somite stage S-5. Our data indicates that the adaxial cell layer undergoes a large-scale tissue shear across the dorsal and ventral regions. This forms a chevron-like pattern which is sharpest at stage S-2 to S-1, before relaxing during somite segmentation. Further analysis shows that this tissue shear is mediated by directional cell intercalations of adaxial cells. These intercalations appear to be dependent on the apical constriction of adaxial cells via apically localized non-muscle myosin IIA in a Shh dependent manner. Our results show that formation of somite segments requires orientated cell rearrangements and spatially distinct cell morphological changes mediated, at least in part, by Shh signalling. This highlights a previous unknown link between Shh and cell mechanical properties in the PSM and provides insight into how cell boundaries form within tissues undergoing complex morphological changes.

## Results

### The shape of future somite boundaries of adaxial cells and lateral somitic cells are distinct within the PSM

After segmentation, each somite undergoes a break in symmetry from a cuboidal shape into a chevron-like morphology orientated along the AP-axis (Figure S1A-B) (Rost et al., 2014; Tlili et al., 2018). Interestingly, deformation of the somite is not uniform, with the more medial layers undergoing larger scale changes earlier in somitogenesis. We first asked whether symmetry breaking also occurs within the PSM and, if so, whether there is spatial inhomogeneity.

We utilized back-tracking of single cells from S3 to S-2 to identify the future somite boundaries in both adaxial cells and lateral somitic cells within the PSM (Figure 1A-C). We noticed that the future boundary of adaxial cells displays a chevron shape at somite stage S-2 (Figure 1A(i)). In contrast, the future boundary of lateral somitic cells is roughly straight across the dorsal-ventral (DV)-axis at S-2, though irregular at the scale of single cells (Figure 1B(i)). During and after somite segmentation, the highly curved border of the adaxial cell population gradually aligns with that of the lateral cells (Figure 1C(iii-vi)). By stage S3, the somite boundaries of adaxial and lateral somitic cells are well aligned (Figure 1A-C). To quantify the temporal-spatial variation between the somite boundaries defined by adaxial cells and lateral cells, we quantified the maximum deviation of the two boundaries along the AP axis (which always emerges near the DV midline) (Figure 1D). At S-2 there is a clear difference in the boundaries, but this rapidly decreases during somite segmentation (Figure 1D’).

**Figure 1.**
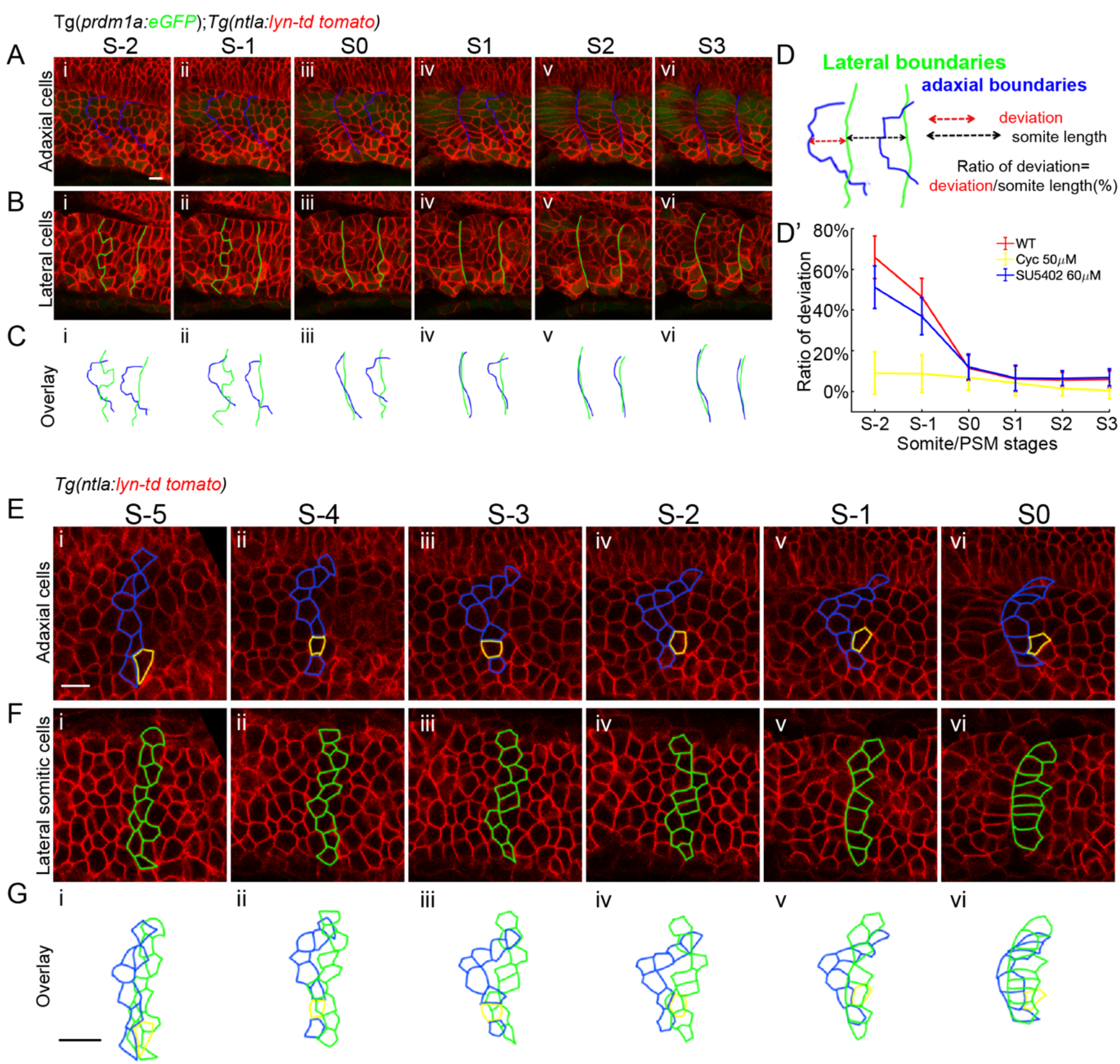
The divergent somite borders of adaxial cells and lateral somitic cells are due to differential cell movement within the PSM. (A-C) Somite boundaries at the adaxial cells and lateral somitic cells from stage S-2 to S3: (A) adaxial cells; (B) lateral somitic cells; and (C) overlay of the two different boundaries. Anterior to the left and dorsal to the top in all parasagittal views unless otherwise stated. (D) AP deviation of somite boundaries of the two cell types measured at the DV midline. Deviation is defined as the ratio of the midline deviation to the AP length of the corresponding somite. (D’) Changes of ratio of AP deviation to the somite length in wild type embryos (n_Somites_=7, n_Embryos_=5), embryos under cyclopamine treatment at 50 μM (n_Somites_=7, n_Embryos_=5) or SU5402 treatment at 60 μM (n_Somites_=6, n_Embryos_=6). (E-G) Cell tracking of adaxial cells (E) and lateral somitic cells (F) which are located at the anterior somite border at somite stage S0 (E(vi) and F(vi)). Cells are traced back to somite stage S-5 (E(i) and F(i)). (G) Overlay of the boundary adaxial cells and boundary lateral somitic cells from (E-F). Images are taken at somites 18-20. Scale bars 10 μm.

Shh signaling is required for the differentiation of adaxial cells into slow muscles but not the initial induction of adaxial cells (Hirsinger et al., 2004). To test whether Shh plays a role in morphogenesis in the PSM we treated embryos with cyclopamine, a small molecular inhibitor of the downstream activator Smoothened (Smo) (Chen et al., 2002). Under inhibition of Shh activity, both adaxial cells and lateral somitic cells are segmented in straight lines at similar locations (Figure 1D’ and S1C-E). In contrast, inhibition of FGF signaling with SU5402 - under which the positions of somite boundaries are altered (Dubrulle et al., 2001) - leaves the difference in boundaries between adaxial and lateral somitic cells largely intact (Figure 1D’ and S2A-D). Therefore, Shh signaling appears to be important in determining the distinct boundaries between cell layers and this process is uncoupled from the boundary specification itself.

### Tissue shear shapes the future somite boundary within the adaxial cell layer

To identify the origin of the distinct future somite boundaries within the PSM, we back-tracked both adaxial and lateral somitic cells from stage S0 to stage S-5, when the future somite boundary is determined by FGF and its downstream factors (Figure S3A-B) (Akiyama et al., 2014). Consistent with previous work, we find that the initial border of both adaxial and lateral somitic cells are straight in the DV axis, and are located nearby to each other at stage S-5 (Figure 1E(i)-G(i)).

From stages S-5 to S-1, the adaxial cell population is reshaped, with changes in connectivity and cell packing (Figure 1E). During this tissue rearrangement, the dorsal and ventral adaxial cells move posteriorly relative to the midline adaxial cells (Figure 1E, Movie 1), thus generating a tissue shear. During this process the epithelial-like adaxial cell layer remains congruent, with no clear separations between cells, though they display highly plastic cell-cell adhesions allowing tissue remodelling. In contrast, we observed less obvious changes in the relative positions of lateral somitic cells from stages S-5 to S-2. During stage S-2 to S0, these cells aligned more with each other, forming a more continuous boundary with a slight U shape forming at the lateral border (Figure 1F). These differential cell rearrangements explain the differing patterns of somite segmentation between the cell populations (Figure 1G), even though the borders of each cell type are determined at the same time and position. Effectively, the adaxial cell population appears to undergo shearing, whereas the lateral somitic cells remain largely static relative to each other.

To further explore the dynamics of this effective tissue shear and its potential role in the tissue shape formation, we quantified the angle of the boundary adaxial cells (BACs) located next to the anterior border of each future somite (Figure 2A). Three phases of boundary shape formation can be demarcated according to the dynamic profile of the angle (Figure 2B). The first phase (S-5 to S-1.5) corresponds to the tissue shear, in which the shear flow initiates the formation of a chevron-like shape in the medial BACs. In this phase, the angle of BACs changes rapidly from 183±4° to 86±12° (Figure 2B). During the second phase, from S-1.5 to S1, the tissue appears to undergo remodeling, whereby the borders of the adaxial and lateral somitic cells become increasingly aligned. The angle of the BACs increases from 86±12° to 123±5°, possibly due to the whole-tissue remodeling of the somite during segmentation from the PSM (Figure 2B). In the third phase, the shape of the future myotome (derived from the somite) begins to emerge. During this phase, the angle of the somite boundaries sharpens before stabilizing (Figure S1A-B(iv)) (Tlili et al., 2018). Thus, we see that spatial symmetry breaking occurs noticeably before somite segmentation and has distinct behaviors in different regions of the PSM.

**Figure 2.**
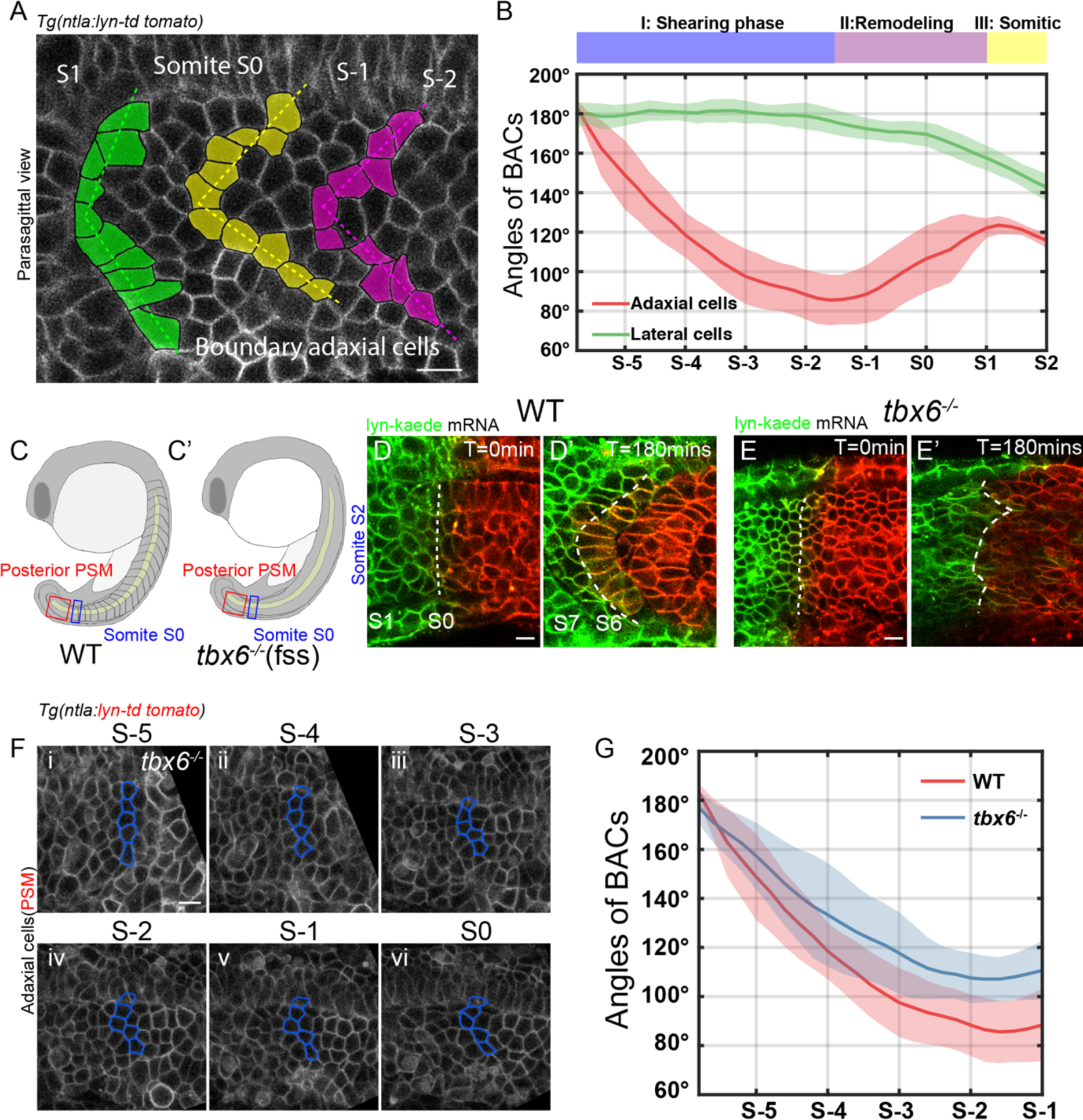
Tissue shear occurs in the adaxial cell layer within the PSM and is independent of pre-existing somites. (A) Adaxial cells located at the anterior somite boundary display a chevron-shaped pattern within the PSM. The angles of the anterior boundary cells are measured at the crossing of lines that are fitted according to the centroids of the dorsal and ventral cells respectively. (B) Angle formed by the boundary adaxial (red) and lateral somitic cells (green) from stage S-5.5 to S2 (n_Somites_=5, n_Embryos_=4). Violet, pink and yellow bars denote the shearing phase, remodeling phase and somitic phases respectively as discussed in the text. (C) Schematic diagram of wild type embryos (C) and *tbx6^-/-^* mutants (C’) at 18-somite stage. (D-E’) Photo-conversion of a photo-convertible protein Kaede in wild-type embryos (D) and *tbx6^-/-^* mutants (E). Photo-conversion was performed at somite S0 in wild type embryos (D) or equivalent region of *tbx6^-/-^* mutants (E) (blue rectangle region in C and C’). (D’-E’) 180 mins after the photo-conversion shown in D-E. Dashed white lines label the borders between the converted and unconverted somitic cells. (F) Cell tracking of adaxial cells in *tbx6^-/-^* mutants from stage S-5 to S0 (red rectangle region in C and C’). Blue color denotes adaxial cells located at the similar AP positions at stage S-5. (G) Profiles of the angles of the boundary adaxial cells in wild-type embryos (red) and *tbx6^-/-^* mutants (blue). All scale bars 10 μm.

The distinct cell fates within the future somite segment are Shh dependent. Therefore, we next asked whether the tissue deformations in the PSM are dependent on Shh. We treated embryos with cyclopamine and repeated the cell backtracking procedure as above to determine cell position within specific future somites. Intriguingly, we found that the tissue shear is largely absent under cyclopamine treatment. This suggests that Shh signaling is important in shaping the future somite (Figure S3C-E), and we return to this later.

### Tissue shear is independent of pre-existing somite boundaries

We next explored the possible mechanisms for induction of the tissue shear within the PSM adaxial cell population. One possible explanation is that the chevron-like shape of older somites acts as a template for more posterior tissues. However, the angle of boundary adaxial cells at stage of S-1.5 (∼90°) is even smaller than the angle of a typical mature somite (∼110°) (Figure 2A-B and S1B(iv)). Thus, the shape of older somite segments cannot account for the tissue deformation of adaxial cells in the PSM. To test this further, we took advantage of somite boundary disruption in *tbx6^-/-^* mutants (Figure 2C) (Nikaido et al., 2002). Using lyn-kaede to demarcate regions of the somite in wild-type (Figure 2D) and *tbx6^-/-^* mutants (Figure 2E) during somitogenesis, we see no tissue deformation in *tbx6^-/-^* mutant somites from S2-S6, suggesting that formation of somite boundaries is necessary for correct morphogenesis of the mature somite. However, the tissue shear of the PSM adaxial cell population still occurs in *tbx6^-/-^* mutants, though it is less stark as compared with wild-type embryos (Figure 2F-G). Therefore, the tissue flow of the adaxial cells appears to be largely independent of the pre-existing chevron-shaped somites and the shear of anterior tissues.

### Adaxial cell rearrangements are polarized within the PSM

Given that the tissue shaping is not due to the pre-existing somites, we next explored the cell rearrangements within the PSM itself as a potential cause of the tissue morphogenesis. During somite stages from 11 hpf to 24 hpf, tailbud progenitor cells continuously exit and differentiate into elements of the growing body (Lawton et al., 2013). Precursors of the adaxial cells are produced by the posterior wall progenitor cells (PWPCs, also referred as neuromesodermal progenitors), which are bipotential neural and mesodermal progenitors located at the posterior end of the tail bud (Gouti et al., 2014; Row et al., 2016). The earliest adaxial cells can be identified at somite S-6 or even earlier in the posterior PSM (Methods and Figure S4A). These early adaxial cells display irregular cell shape but with extension along the DV direction (Figure S4A-B). They converge towards the DV midline and join the pseudo-epithelial array, with their aspect ratio decreasing towards one from somite stage S-6 to S-3 (Figure S4A-B). Interestingly, the aspect ratio of adaxial cells changes in a similar manner in Shh inhibited embryos (using cyclopamine) compared with wild type embryos (Figure S4B-C). This is consistent with previous work, which showed that the initial induction of the adaxial cells is independent of Shh signalling (Hirsinger et al., 2004).

We quantified the cell rearrangements of the adaxial cell within the PSM (Figure 3A). We noticed that adaxial cells undergo polarized cell intercalations along the DV axis. Most cell intercalations were orientated towards the DV-axis (Figure 3B). Here, we denote the two cells that share the edge before T1 transitions as b1 (anterior) and b2 (posterior) and cells that are approaching to each other as a1 (dorsal) and a2 (ventral) (Figure 3B’). The directional T1 transition along the DV direction induces convergence of cells along the DV direction by pulling dorsal cell a1 and ventral cell a2 together. Meanwhile, cells are further compacted along the AP direction by the orientated separation of cells b1 and b2 (Figure 3A and 3B’).

**Figure 3.**
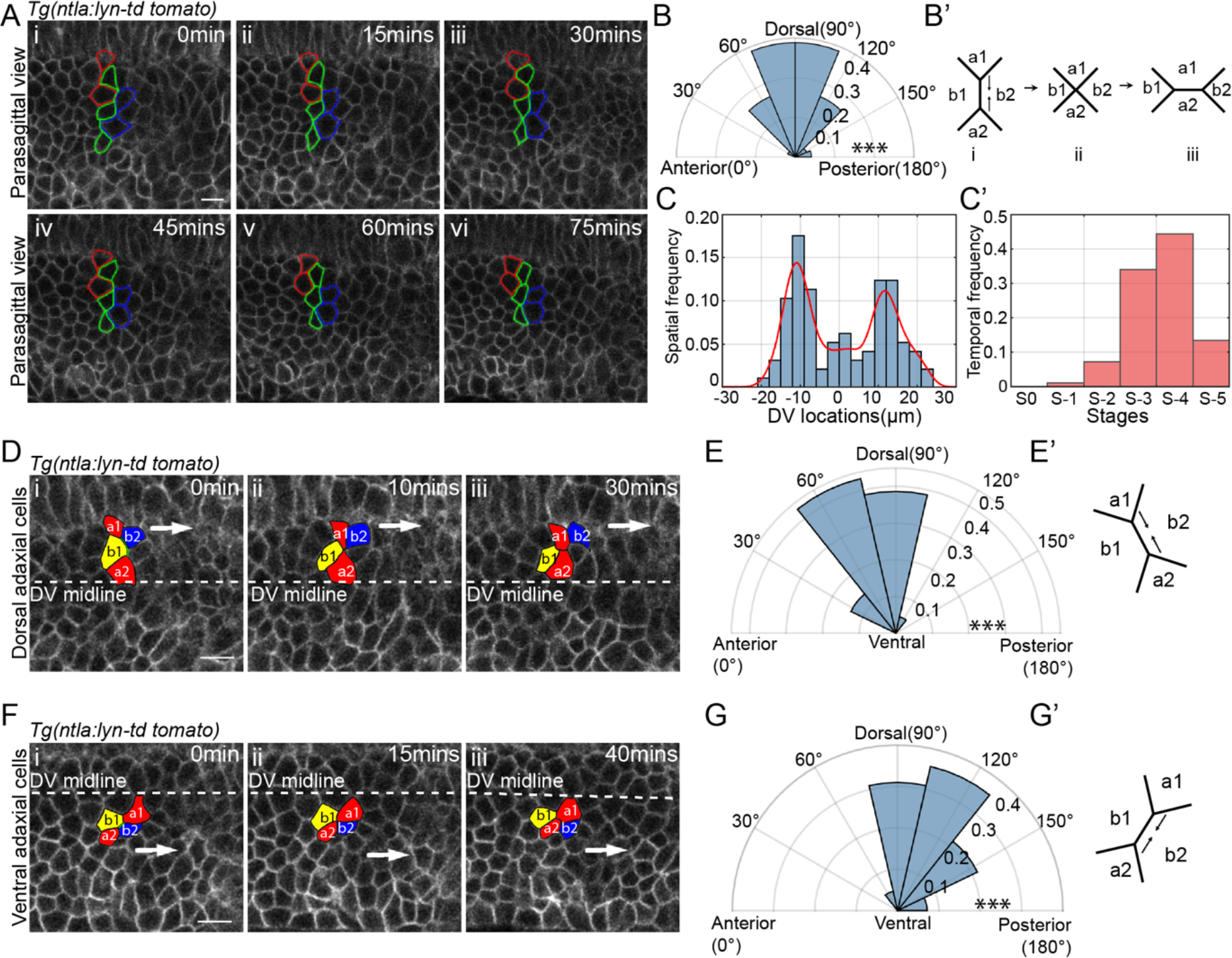
Directional cell intercalations contribute to the tissue shear of adaxial cells. (A) Time lapse of intercalations in wild type embryos of adaxial cells from stage S-5 (i) to S-3 (vi). Red, green and blue denote pairs of converging cells. (B) Distribution of the orientations of adaxial cell intercalations (n_Intercalations_=97, n_Embryos_=5). The orientations of intercalations are measured according to the direction of the constricting edge shared by cells b1 and b2 before the T1 transition (B’). (B’) Diagram of T1 transition with the converging cells denoted as a1 (dorsal) and a2 (ventral) and separating cells denoted as b1 (anterior) and b2 (posterior). (C-C’) The spatial (C) and temporal (C’) frequency of intercalations of the adaxial cells (n_Intercalations_=97, n_Embryos_=5). Red line in (C) denotes to the kernel density estimation of the spatial probability of intercalations along the DV axis with bandwidth of 2. (D-G) Time lapse of adaxial cell intercalations occurring at dorsal (D) and ventral (F) regions. Long white arrows denote the directional of shear of dorsal and ventral cells respectively. Distribution of adaxial cell intercalation orientation at the dorsal (regions 4 μm dorsally above the midline) (n_Intercalations_=44, n_Embryos_=5) (E) and ventral (regions 4 μm ventrally below the midline) (n_Intercalations_=42, n_Embryos_=5) (G) regions. Diagrams of the dorsal intercalations (E’’) with an anterior-ward fraction and ventral intercalations (G’’) with a posterior-ward fraction. Images are taken at somites 18-20. Scale bars 10 μm. ***p < 0.001, Rayleigh test was performed to test the non-uniformity of the directions of adaxial cell intercalations.

We quantified the temporal and spatial occurrence of adaxial cell intercalation throughout the PSM from stage S-5 to S-1. Interestingly, the spatial occurrence of adaxial cell intercalations display a bimodal distribution, with the highest frequency at the dorsal (+10 μm) and ventral sides (−10 μm) and lowest frequency at the midline (Figure 3C). Consistent with the jamming transition model recently reported (Mongera et al., 2018), the intercalations of adaxial cells primarily occurs in the posterior PSM from S-3 to S-5 (Figure 3C’). Finally, returning to our observation above regarding the dependency on Shh for tissue shear, we tested whether these intercalations were Shh dependent. We find that, in the absence of Shh activity, the adaxial cells remain largely static with few cell rearrangements (Figure S4D). This suggests that cell morphological changes regulated by Shh are crucial for morphogenesis of the tissues within the PSM.

### Directional cell intercalations induce tissue shear of adaxial cells

We have seen that there is spatial symmetry breaking in the adaxial cell layer induced by a tissue shear and that cells within the PSM have spatially varying rates of cell rearrangements. Can these directional cell intercalations drive the observed tissue shear? To answer this, we carefully examined the changes in cell shape and cell connectivity throughout the polarized cell rearrangements. First, we compared the relative cell extension of cells b1 and b2 (as defined above) along the AP axis while undergoing a T1 transition (Figure S5A). The relative positions of the cells b1 and b2 along the AP axis are measured relative to the centroids of cells a1 and a2 at the time of four-way junction (denoted t_2_) and after the separation of cells b1 and b2 (denoted t_3_) (Figure S5A). By comparing the relative cell movements, we found that the posterior cell b2 extends a longer distance compared to the anterior cell a1 (Figure S5B). Thus, the rearrangements along the DV axis lead to a relatively larger extension in the posterior direction, as compared to the anterior elongation. Combined with the spatially bimodal distribution of adaxial cell rearrangements (Figure 3C), the dorsal and ventral located cells extend further posteriorly compared with cells at the midline and with higher frequency of rearrangements.

We also noticed a proportion of rearrangements orientated along the AP-axis. To look further into the possible contribution of these rearrangements in shaping tissues, we separately quantified such events at distinct spatial locations (dorsal, ventral, midline). A significant proportion of the dorsally located rearrangements displayed an anterior-ward orientation, whilst most of the ventral rearrangements displayed a posterior-ward orientation (Figure 3D-E). Such orientated rearrangements contribute to the tissue shear flow of adaxial cells by rearranging the connectivity of the intercalating cells by generating a posterior-ward tissue shear at the dorsal and ventral sides of the adaxial cell layer (Figure 3F-G). The delicate balance in the orientation of these different rearrangements induces the tissue shear flow and symmetry breaking within the adaxial cell layer in the PSM.

### Apical constriction of adaxial cells drives tissue shear in a Shh dependent manner

We have shown above that directional cell intercalations induce tissue shear of adaxial cells. To study the basis of the directional cell intercalations further, we traced every adaxial cell within an emerging somite from stage S-5 to S0 and recorded their shape changes in the parasagittal plane at the apical-basal midplane (Figure 4A). In wild type embryos, adaxial cells converged towards the dorsal-ventral midline with the average cell cross-sectional area at the apical-basal midplane reducing from 63±12 to 43±9 μm^2^ from stage S-5 to S-0 (Figure 4A-B). In contrast, though displaying similar cell size at S-5 compared with wild type embryos, adaxial cells in *smo^-/-^* mutants (Barresi et al., 2000; Wolff et al., 2003) or wild type embryos under cyclopamine treatment show little change in cell size (Figure 4B). Interestingly, the reduction of cell size is not uniform in untreated wild type embryos. The dorsal and ventral adaxial cells display larger reductions in size compared with midline cells in wild type embryos (Figure 4C).

**Figure 4.**
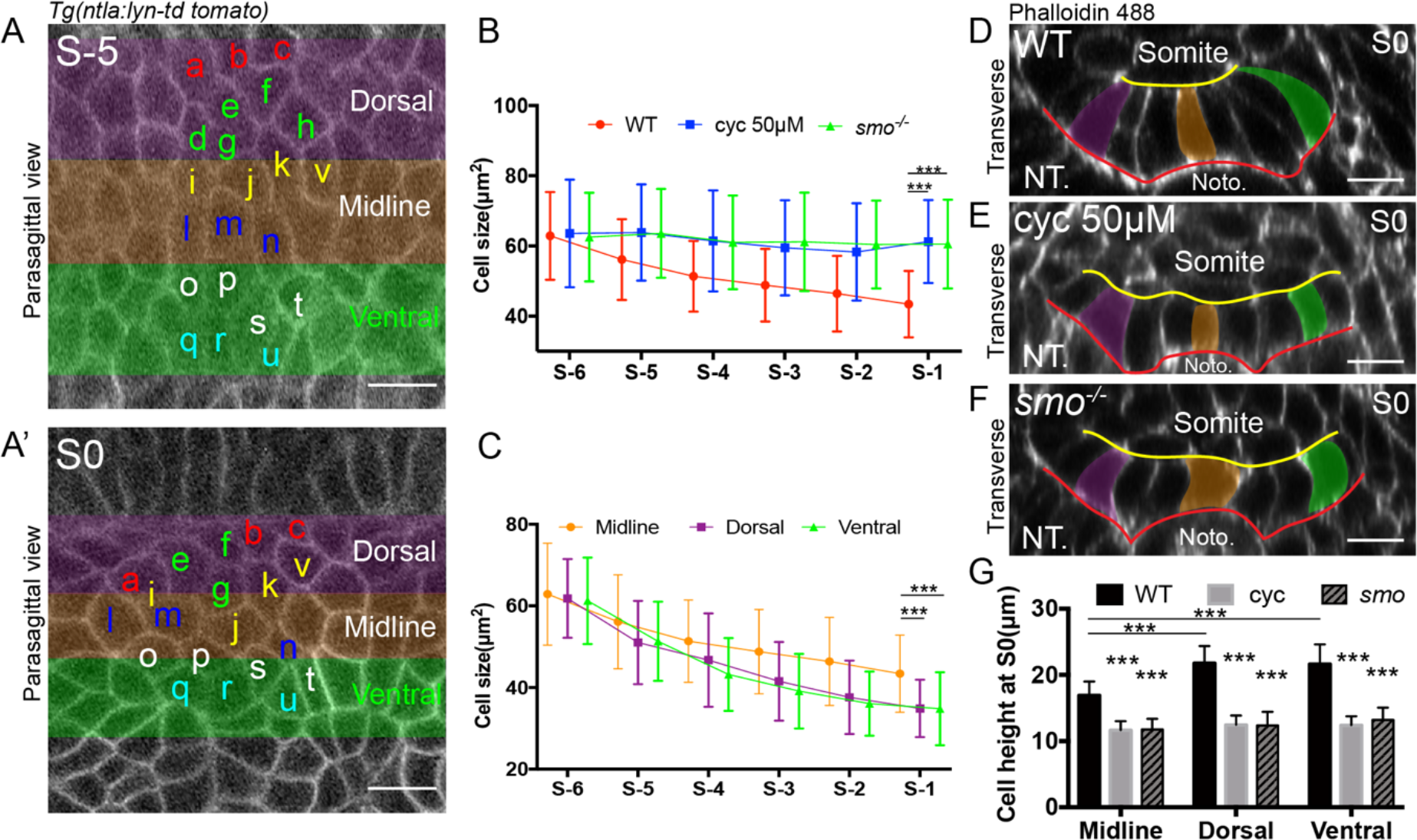
Polarized morphogenesis of adaxial cells is Shh dependent. (A) Cell tracking of adaxial cells in a wild type embryo from a future somite at stage S-5 (A) to stage S0 (A’). Shaded regions denote dorsal, midline and ventral regions respectively. (B) Adaxial cell cross-sectional area measured at the apical-basal midplane in different stages (n_Cells_=43, n_Embryos_=7), embryos under 50 μM cyclopamine treatment (n_Cells_=26, n_Embryos_=4) and *smo^-/-^* mutants (n_Cells_=35, n_Embryos_=6). (C) Adaxial cell cross-sectional area measured at the apical-basal midplane in different stages for cells located at the dorsal (n_Cells_=32, n_Embryos_=7), midline (n_Cells_=43, n_Embryos_=7) and ventral regions (n_Cells_=32, n_Embryos_=7) in wild type embryos. (D-F) Reconstructed transverse sections at somite stage S0 in wild type embryos (D), embryos under 50 μM cyclopamine treatment (E) and *smo^-/-^* mutants (F) with dorsal to the left and lateral to the top. Violet, orange and green denote dorsal, midline and ventral located cells respectively. Scale bars 10 μm. ***p < 0.001, Student’s t test.

The adaxial cells constrict at their apical surface towards the DV midline in the anterior PSM of wild type embryos (Figure 4D, Movie 2) (Daggett et al., 2007). In contrast, we observed little apical constriction of adaxial cells in the absence of Shh activity (Figure 4E-F, Movie 3). Adaxial cells at somite S0 remain cuboidal in *smo^-/-^* mutants or in wild type embryos under cyclopamine treatment (Figure 4G). By measuring the cell height of adaxial cells from somite S-6 to S-0 in the coronal optical planes (Figure 5A-C), we noticed that the cell height gradually increases from S-6 to S0 in wild type embryos (Figure 5D). The ML extension of adaxial cells is significantly compromised under inhibition of Shh signalling (Figure 5B-D). Consistent with the differential changes of cell size, the dorsally- and ventrally-located cells are significantly taller than the midline cells in wild type embryos, and display stronger apical cell constriction (Figure 4D and 4G). These morphological variations are significantly compromised in the absence of Shh activity, which also displays a loss of apical constriction in the adaxial cell layer (Figure 4E-G).

**Figure 5.**
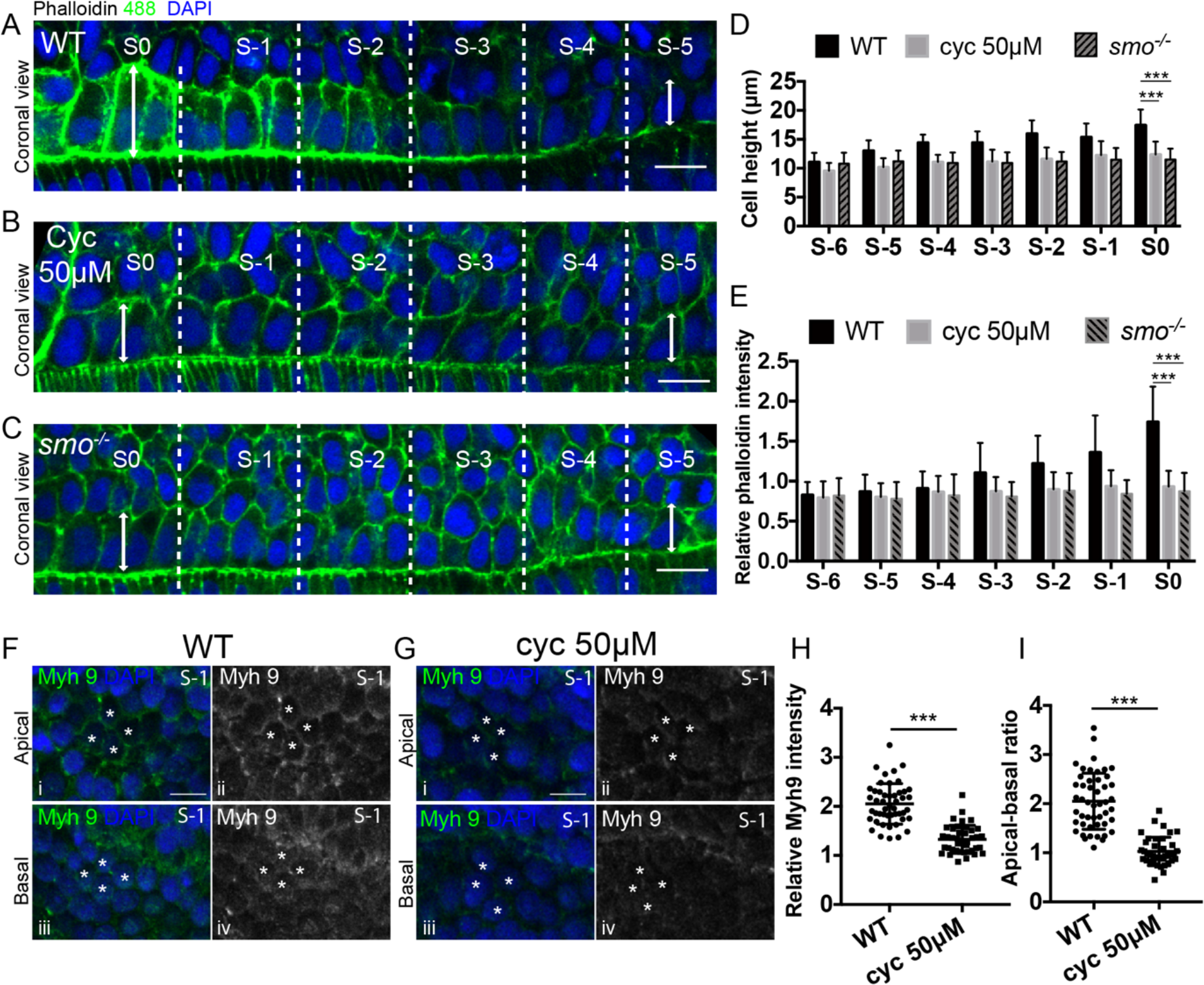
Shh dependent spatial localization of Actin and Myh9 is necessary for polarized cell constriction. (A-C) Coronal view of the PSM from somite S-5 to S0 in wild-type embryos (A), embryos under 50 μM cyclopamine treatment (B) and *smo^-/-^* mutants (C) with anterior to the left and lateral to the top. Cell membranes are labelled with Phalloidin 488 and cell nuclei with DAPI. (D) Cell height (ML length) of adaxial cells from somite S-6 to S0 in fixed wild-type embryos (n_Cells_=152, n_Embryos_=7), embryos under 50 μM cyclopamine treatment (n_Cells_=166, n_Embryos_=7) and *smo^-/-^* mutants (n_Cells_=161, n_Embryos_=7). (E) Relative Phalloidin intensity at the lateral surface of adaxial cells from somite S-6 to S0 in fixed wild-type embryos (n_Cells_=350, n_Embryos_=5), embryos under 50 μM cyclopamine treatment (n_Cells_=350, n_Embryos_=5) and *smo^-/-^* mutants (n_Cells_=350, n_Embryos_=5). Phalloidin intensity normalized according to the average intensity at the membrane of lateral cells in the same embryo. (F-G) localization of Myh9 at the apical (i-ii) and basal (iii-iv) sides of the lateral surface of adaxial cell in wild type embryos (F) and embryos under cyclopamine treatment at 50 μM (G). White asterisks label adaxial cells at stage S-1. (H) Relative intensity of Myh9 at the apical lateral side of adaxial cells in wild type embryos (n_Cells_=48, n_Embryos_=6) and embryos under cyclopamine treatment at 50 μM (n_Cells_=41, n_Embryos_=6). The Myh9 intensity was normalized according to the average intensity of Mhy9 in the lateral region of the same embryo. (I) Distribution of the ratios of Myh9 intensities between apical and basal sides of the lateral surface in wild type embryos (n_Cells_=48, n_Embryos_=6) and embryos under cyclopamine treatment at 50 μM (n_Cells_=41, n_Embryos_=6). Images are taken at 18-somite stage. Scale bars 10 μm.***p < 0.001, Student’s t test.

### Myosin-II distribution is polarized in adaxial cells within the PSM

Directional cell intercalations, mediated through apical constriction, induce tissue shear of adaxial cells. We next probed the molecular processes underlying constriction of adaxial cells. Interestingly, we observed strong enrichment of actin at the apical side of anterior adaxial cells (Figure 5A and 5E). In contrast, the intensity of actin appears largely unchanged from somite S-5 to S0 in the absence of Shh activity (Figure 5B-C and 5E). Given its known role in driving apical constriction (Martin et al., 2009), we investigated the cellular distribution of non-muscle myosin II A (Myh 9) in adaxial cells. Myh 9 appears to localize at the apical side of the lateral surface in wild-type embryos instead of the basal side (Figure 5F). This observation is consistent with the significant constriction of adaxial cells at the apical surface. Under inhibition of Shh signalling, the apical intensity of Myh9 is significantly reduced and the polarized distribution of Myh 9 is lost, with significant variability in Myh 9 localization (Figure 5H-I and Figure S6A). To test further the role of cell contractility in shaping the tissue shear flow of adaxial cells, we utilized blebbistatin to inhibit non-muscle myosin II. Under the inhibition of non-muscle myosin II, the adaxial cell layer remains largely static, displaying little convergence-extension or tissue shear (Figure S6B). Shh appears to control the morphogenesis and migration of adaxial cells within the PSM by inducing apical constrictions of adaxial cells towards the DV midline.

## Discussion

There has been significant work on characterizing the temporal and spatial control of somite boundary determination in vertebrates (Gibb et al., 2010; Pourquié, 2011; Saga and Takeda, 2001). In this study, we demonstrated that future muscle fibers within the PSM undergo shearing, which appears to be required for robust formation of the somite. This morphological process is not a direct result of the initial boundary formation via FGF signaling or a passive response to the shape of pre-existing somites, but instead it is actively shaped by cell morphogenesis and migration after initial boundary formation of the future somite within the adaxial cell population. After the future somite boundary determination at S-5, the adaxial cells converge towards the DV midline with extensive directional intercalations occurring in dorsal and ventral regions (Figure 6A). There is a spatially-dependent fine-tuning of cellular rearrangements, which mediate the tissue shear of dorsal and ventral adaxial cells towards the posterior direction and subsequently results in symmetry breaking of the adaxial cell layer within the PSM (Figure 6B-C).

**Figure 6.**
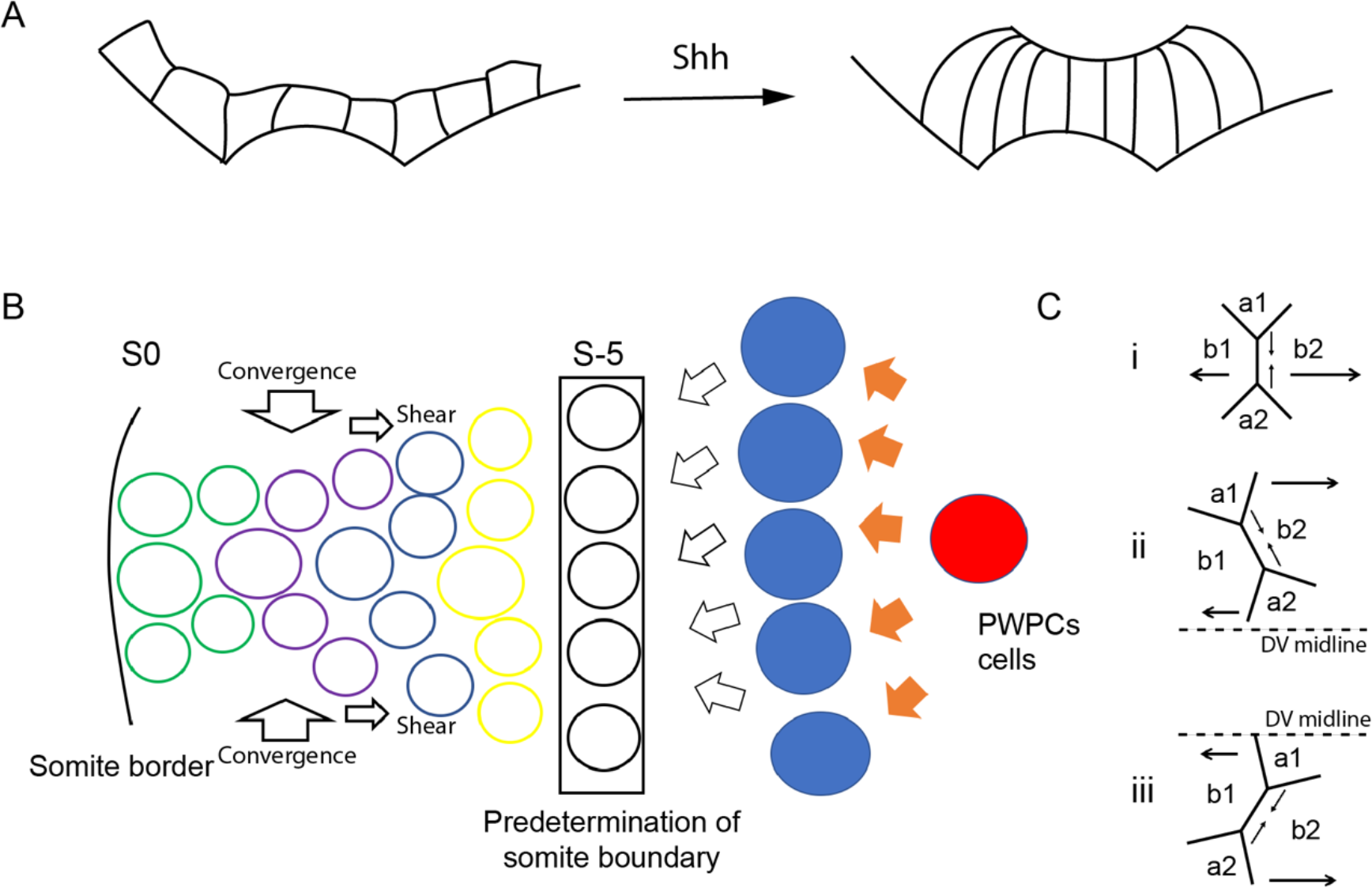
Schematic of adaxial cell morphogenesis induced tissue shear. (A) Adaxial cells undergo directional convergence within the PSM, with their apical side constricting towards the DV midline. (B) Future somite boundaries are determined at somite S-5. The subsequent directional convergence leads to shear of adaxial cells with asymmetric (C (i)) and directional C (ii, iii) cell intercalations shaping the cell morphology and connectivity.

Our results indicate that adaxial cells in the PSM undergo convergence towards the DV midline in the parasagittal planes. This observation is distinct from the convergence and extension of the paraxial progenitor cells that occur along the medial-lateral direction during vertebrate gastrulation. Such movements during gastrulation arrange adaxial cell progenitors near the notochord (Myers et al., 2002; Yin and Solnica-Krezel, 2007). The convergence of adaxial cells at the PSM involves complicated cell morphogenesis and directional cell rearrangements. It has been shown that cell rearrangements are driven by actomyosin-mediated constriction in other systems (Bertet et al., 2004; Rozbicki et al., 2015; Skoglund et al., 2008; Walck-Shannon and Hardin, 2014). During the rearrangements of adaxial cells, cells constrict in their size and extend along the ML direction. Notably, adaxial cells apically constrict towards the DV midline, induced, at least in part, by polarized non-muscle myosin II. With the constricted apical and curved basal surfaces, the dorsal and ventral positioned adaxial cells constrict more than cells positioned near the midline. This appears to lead to higher rearrangement frequencies at the dorsal and ventral sides compared to the midline region.

Geometric constraints and tension are implicated in mediating cell shape changes and cell morphogenesis. Correspondingly, developmental programs must accommodate mechanical tissue forces in order to maintain robust morphogenesis (Shawky and Davidson, 2015; Zhou et al., 2015). Consistent with the key role of cortical F-actin density in mechanically regulating convergence and extension (Shawky et al., 2018), we observed increased intensity of actin during the convergence of adaxial cells at the PSM. We have previously identified apical-to-basal neighbor exchanges in the highly curved anterior of the early *Drosophila* embryo (Rupprecht et al., 2017). Such apical-to-basal neighbor exchanges release high tension due to dense cell packing in regions of high curvature. Interestingly, we observed similar apical-to-basal neighbor exchanges in the adaxial cell layer (Movie 2). This suggests that both spatial and temporal cell rearrangements can occur during morphogenesis even in tissues without high curvature, as has also been observed in *Drosophila* germ-band elongation (Sun et al., 2017). Further, the directional cell intercalation of adaxial cells, which primarily occur along the DV direction but also with a small bias in the AP direction, may facilitate the release of high tension induced within the dorsal and ventral regions by cell shape change. One unanswered question here is why the shear only occurs toward the posterior direction. The unidirectional flow of adaxial cells can be explained by the differential physical constraints at the anterior/posterior ends of the adaxial cell layer. Elongated slow muscles and somite borders located at the anterior side of the adaxial cell layer are much stiffer than the epithelial cells (Butcher et al., 2009; Cox and Erler, 2011). In contrast, the progenitor cells located at the posterior side are more motile and flexible (Mongera et al., 2018; Row et al., 2016). Under the differential constraints, the cells tend to extend towards the posterior direction. The observation that cells tend to extend towards the posterior during T1 transitions in the PSM further support this idea. During *Drosophila* gastrulation, geometrical and mechanical constraints are cues in orienting the actomyosin meshwork and tension during ventral furrow formation (Chanet et al., 2017). Further, it has been observed cells undergo orientated divisions to limit tension anisotropy in epithelial spreading during zebrafish epiboly, suggesting geometric constraints play critical roles in tissue morphogenesis (Campinho et al., 2013).

Shh signaling is implicated in the differentiation of adaxial cells into slow muscles and the further differentiation of them into muscle pioneers and superficial slow muscles (Devoto et al., 1996; Henry and Amacher, 2004). The morphogenesis of slow muscles occurs soon after somite segmentation, whilst the adaxial cells start to display Shh activity in the posterior PSM (Daggett et al., 2007; Lewis et al., 1999). In this study, we have revealed that Shh plays a role in the PSM in controlling cell morphogenesis and migration of adaxial cells. Our data suggests that Shh is required for the apical constriction of adaxial cells, possibly through recruiting Myh9 to the apical side of the cells. This result is consistent with observations in the *Drosophila* eye imaginal disc, where Hh induces cell ingress by localized recruitment of Myosin II (Corrigall et al., 2007). Combined, this suggests that the Hedgehog pathway guides cellular mechanics and is hence important in tissue morphogenesis as well as cell fate determination.

Interestingly, we noticed that the apical side of adaxial cells tends to constrict towards the DV midline, where it is reported the highest levels of Shh are present. Thus, it is possible Shh signaling is involved in the establishment of polarity along the DV axis. However, we also noticed that cells not in direct physical contact with the notochord tend to have even more significant shape changes compared with the midline cells. Such cells display little Shh activity in the somite and do not commit to a slow muscle fate in the somite (Yin et al., 2018). Hence, it is intriguing to speculate that Shh is not directly involved in the morphogenesis of these distal cells. Instead, active reconfiguration of cell-cell contacts might play a key role in mediating the morphogenesis of adaxial cells. There is also possible crosstalk between Shh and Eph/Ephrin Signaling, as suggested from work on ventral spinal cord patterning (Laussu et al., 2017). Further study is required to characterize the different possibilities and to elucidate the pathway between Shh signaling and cell mechanics.

One of the most striking patterns of zebrafish muscle segments is their distinctive chevron shape, which is believed to be optimal for the maximum muscle contraction during swimming by depositing muscle fibre in a helical manner along each segment (Wedekind, 2007). The chevron shape of muscle segments emerges during somite morphogenesis soon after somite segmentation. There have been a number of models proposed to explain the emergence of this shape (van Raamsdonk et al., 1974, 1979; Rost et al., 2014; Tlili et al., 2018; Wedekind, 2007). It is intriguing that the shape of the adaxial cell layer at S-2 mirrors closely the final shape of the mature muscle segments. This suggests a possible role in pre-printing the chevron shape, though significant further work is required to test this hypothesis.

## Acknowledgements

We thank the IMCB fish facilities and particularly the lab of Philip Ingham for support. We thank Philip Ingham, Olivier Hamant and members of the Saunders lab for critical reading of the manuscript. This work was supported by a National Research Foundation Singapore Fellowship awarded to T.E.S. (NRF2012NRF-NRFF001-094) and a Singapore Ministry of Education Academic Research Fund Tier 3 grant (MOE2016-T3-1-002).

## Author contribution

J.Y. and T.E.S. designed the study. J.Y. performed all experiments. J.Y. analyzed the data with assistance from T.E.S.. J.Y. and T.E.S. wrote the manuscript.

## Methods

### Fish strains and maintenance

Maintenance of adult fish and experimental procedures involving zebrafish embryos were carried out at the Institute of Molecular and Cell Biology (IMCB) Zebrafish Facility (Biopolis, Singapore) certificated by the Agri-Food and Veterinary Authority (AVA) of Singapore. Adult fish were maintained on a 14h light/10h dark cycle at 28°C. Fish lines carrying the following transgenes or mutations were utilized in this study: Tg(*PACprdm1:GFP*)^i106^ (Elworthy et al., 2008), Tg(*ntla:lyn-td tomato*)^sq8^ (Lee et al., 2013), *smo*^b641/b641^(Varga et al., 2001), *tbx6*^ti1/ti1^(Nikaido et al., 2002). Embryos are obtained through natural spawning by crossing male and female adults aged 3-18 months and raised at 28°C.

### Assembly of DNA constructs and RNA Injection

For synthesis of mRNA for live imaging, plasmids pcs2/SP6-lyn-tdTomato and pcs2/SP6-lyn-kaede were assembled by cloning lyn and tdTomato/Kaede into pcs2+ vector through the restriction sites of BamHI/XhoI and XhoI/XbaI respectively. Plasmids were linearized with NotI and subsequently used in the *in vitro* transcription of *lyn-tdTomato and lyn-kaede* with mMESSAGE mMACHINE™ T7 Transcription Kit (Thermo Fisher). Microinjection of the above mRNA was performed at the one cell stage at the concentration of 50 ng/ml.

### Imaging of zebrafish embryos

Zeiss LSM 700 laser scanning confocal microscope was used for the imaging of zebrafish embryos. For live imaging, low concentration of low melting agarose (0.4%) was used to make sure the tailbud can still properly grow during live imaging. Zebrafish embryos at 18-20 somite stage were mounted into 35 mm glass bottom microwell dishes (MatTek) with 0.4% low melting agarose (MO BIO laboratories) in E3 embryo medium containing 180mg/ml tricaine (MS-222, Sigma-Aldrich). After the agarose solidified, embryos were imaged using a 40x oil-immersion objective at 26 °C with somite segmentation occurs every 40 minutes. For live imaging of the PSM, the time points were 5-10 minutes intervals and the z-stacks were 1-2 μm intervals.

### Drug treatments

The treatment of cyclopamine (ab120392, Abcam) was performed at 50% epiboly stage at a concentration of 50 μM in E3 medium. The dose of drug treatment was based on previous reports (Wolff et al., 2003) to remove almost all of the slow muscles. The treatment of SU5402 was performed 2 hours before live imaging at 14-somite stage at 60 μM, at which concentration the positions of somite borders are altered (Sawada et al., 2001; Yin et al., 2018). Blebbistatin (B0560, Sigma) treatment was performed at 100 μM 2 hours prior to live imaging according to previous study (Urven et al., 2006). During live imaging, same concentrations of inhibitors were maintained.

### Antibody staining, fluorescent *In situ* hybridization and imaging

For the staining of myosin, embryos were fixed by Dent’s fixative (80% methanol, 20% DMSO) overnight at 4°C (Dent, 1989). Anti-Myosin IIA Antibody (M8064, Sigma-aldrich) was used in the immunostaining of Myh9. Embryos were incubated with antibody diluted at 1:250 in 2% BSA overnight at 4°C.

High-resolution *in situ* hybridization of whole-mount zebrafish embryos was performed according to established protocols (Thisse and Thisse, 2008). SIGMAFAST™ Fast Red TR (Sigma) was utilized for generating red fluorescence with alkaline phosphatase on DIG labelled probes. Rabbit Anti-GFP antibody (TP401) was used to stain GFP at a dilution of 1:500. For synthesis of probes of *pea3* and *fgf8a*, PCR was performed to amplify templates with the following primers:

pea3_R: TAATACGACTCACTATAGgctcaccagccaccttttgc

pea3_F ctggaccagcaagtgccttatact

fgf8a_R: TAATACGACTCACTATAGGtcaacgctctcctgagtagcg

fgf8a_F: cacggttgagttatctattccttcacc

Whole mount embryos were imaged using a 40x oil immersive objective under a Zeiss LSM 700 confocal microscope.

### Image processing and statistical analyses

The field of view of the LSM 700 was adjusted according to the drift or the growth of the zebrafish embryos every 1-3 hours. Custom MATLAB codes were utilized to re-align the 3D volumes of images at different time slots. ImageJ were used in the image measurement and 3D reconstruction. The stages of cells in the PSM were identified according to how many cycles of somite segmentation occurred before the corresponding somite epithelized in live imaging. In fixed embryos, the stages of cells at the PSM were deduced according to the distances of the cells relative to somite S1 and the average length of the emerging somite at the AP axis. The area of adaxial cells was measured at the apical-basal midplane. The height (ML length) of the adaxial cells was measured by calculating the length of the fitted curve that pass through the centers of cell at each Z-stack from basal to apical sides.

The zebrafish embryos were divided into different treatment groups without any bias. The embryos in each group were selected for live imaging randomly. At least three independent experiments were performed in each group. No statistical power analysis was used to determine samples size. Systematic randomization was not used. Non-parametric two-sided two sample t-test, paired sample t-test, two sample Kolmogorov-Smirnov test and Rayleigh test were performed in MATLAB.

## Supplementary Information

**Supplementary Figure 1.**
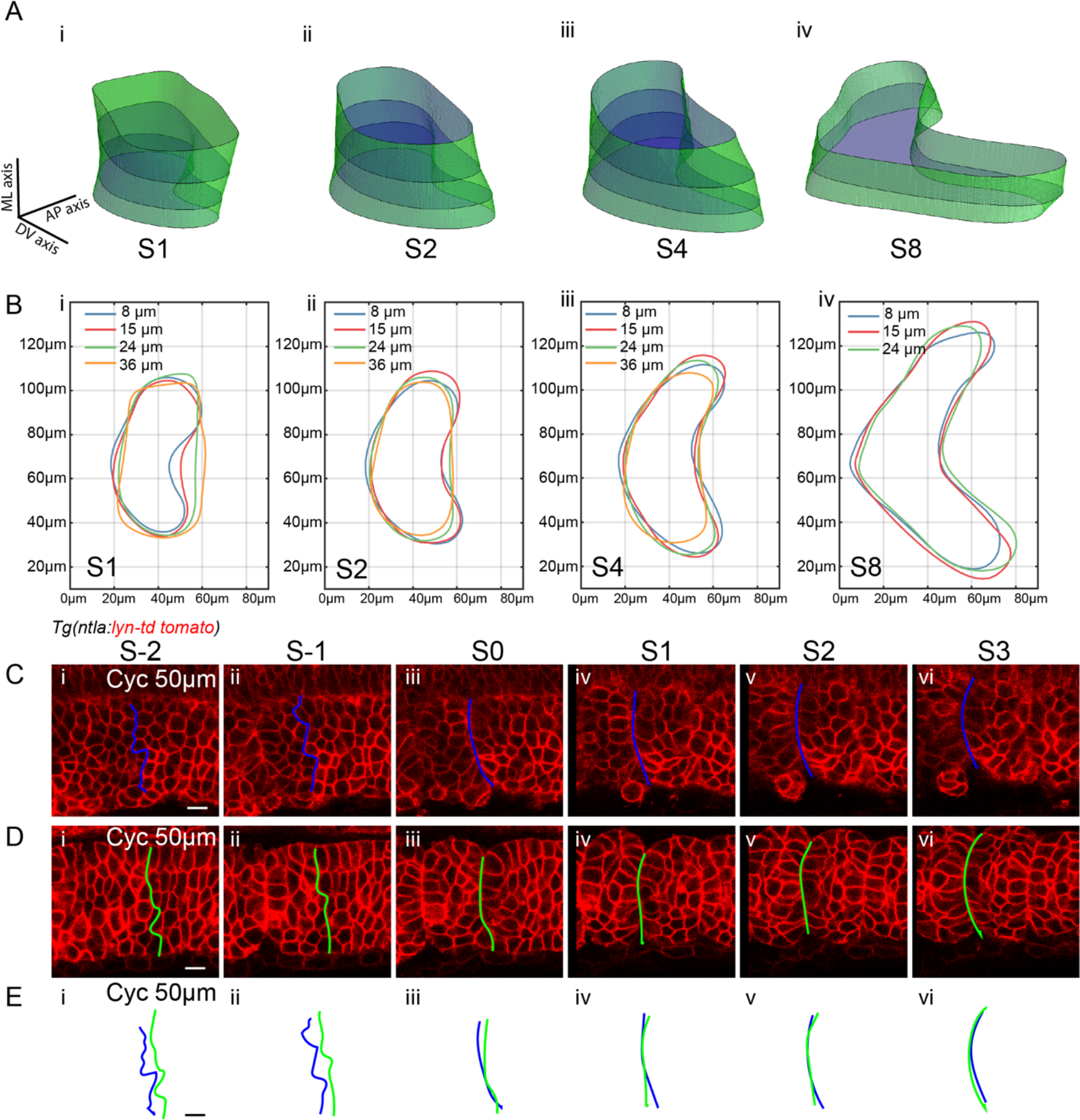
Somite morphogenesis and cell rearrangements under Shh inhibition. (A) 3D somite shapes reconstructed from the contours of somites at parasagittal planes 36, 24, 15 and 8 μm from the notochord. (B) Contours of somites at stage S1, S2, S4 and S8. Colors denote the contours at different ML positions. The contours of somites are taken at somite 18. (C-E) Somite boundaries at the adaxial cells and lateral somitic cells from stage S-2 to S3 under 50 μM cyclopamine treatment: (C) adaxial cells; (D) lateral somitic cells; and (E) overlay of the two different boundaries. Images are taken at somites 18-20. Scale bars 10 μm.

**Supplementary Figure 2.**
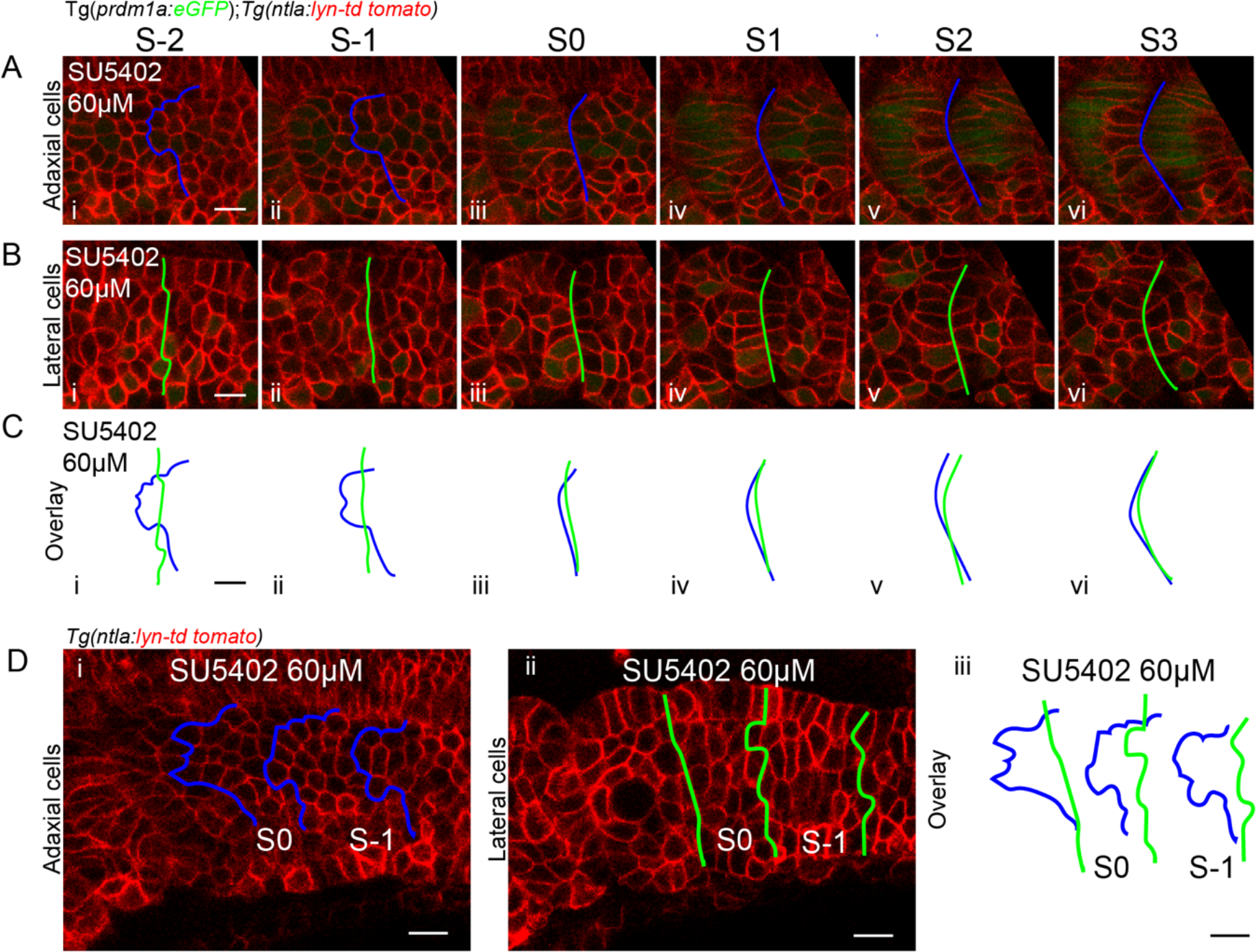
The distinct somite borders at adaxial and lateral somitic cells appear under FGF inhibition. (A-C) Somite boundaries at the adaxial cells and lateral somitic cells from stage S-2 to S3 under 60 μM SU5402 treatment: (A) adaxial cells; (B) lateral somitic cells; and (C) overlay of the two different boundaries. (D) Relatively small (S0) and large (S-1) somite were induced under 60 μM SU5402 treatment, whilst the distinct somite borders at adaxial and lateral somitic cells occur. Images are taken at somites 18-20. Scale bars 10 μm.

**Supplementary Figure 3.**
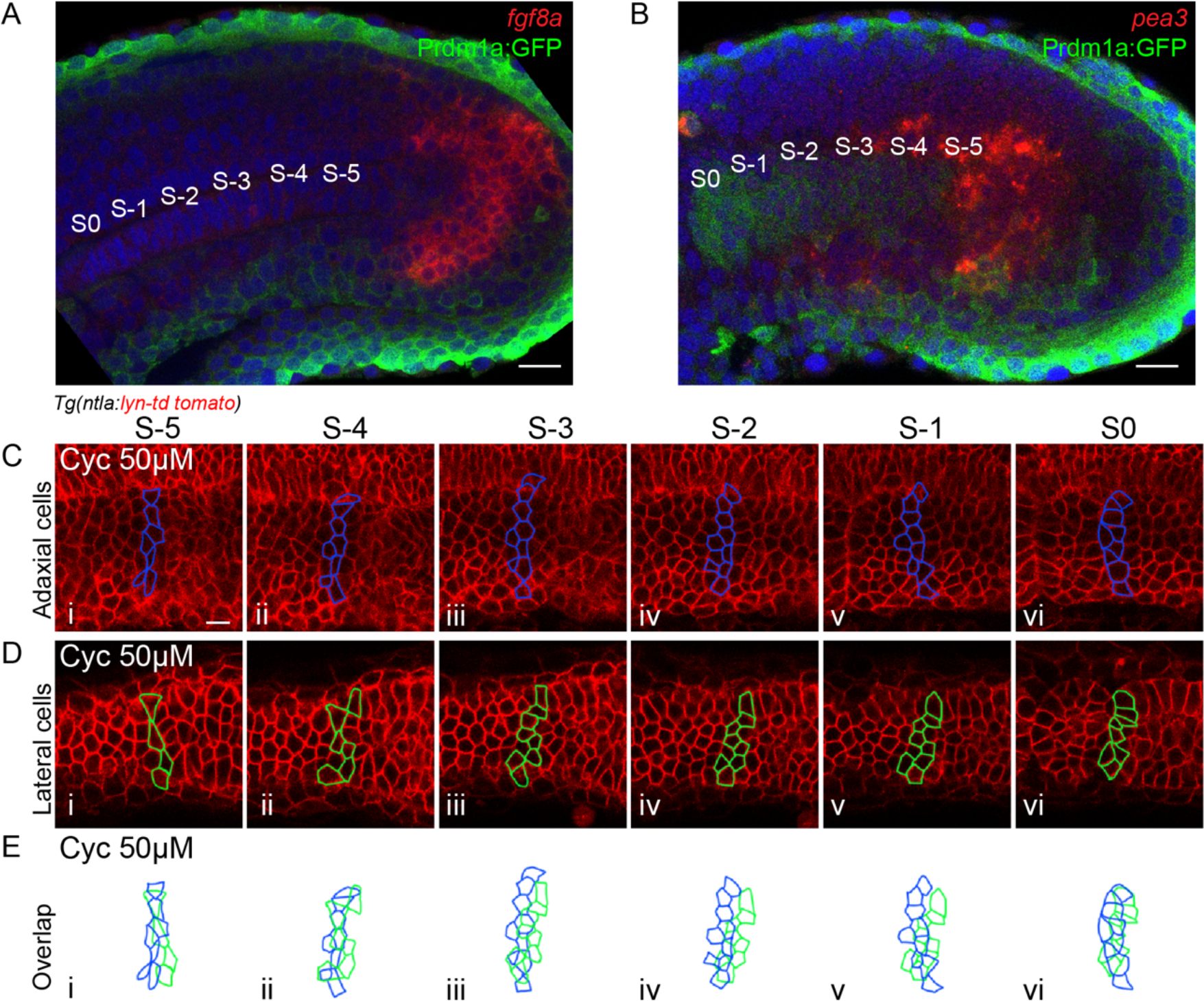
Fluorescent *in situ* of *fgf8a* and *pea3* and cell tracking in the PSM under Shh inhibition. (A-B) Expression of *fgf8a* (A) and a downstream factor of FGF signalling pathway, *pea3* (B), within the tailbud and posterior PSM. Adaxial cells are labelled with Prdm1:GFP. Images are taken from the parasagittal optical planes at the same ML planes of the notochord and adaxial cells respectively. (C-E) Cell tracking identified the adaxial cells (C) and lateral somitic cells (D) located at the anterior somite border at somite stage S0 (C(vi) and D(vi)) in embryos under 50 μM cyclopamine treatment. Cells are traced back to stages as early as somite stage S-5 (C(i) and D(i)). (E) Overlay of the boundary adaxial cells and boundary lateral somitic cells from (C-D). Images are taken at somites 18-20. Scale bars 10 μm.

**Supplementary Figure 4.**
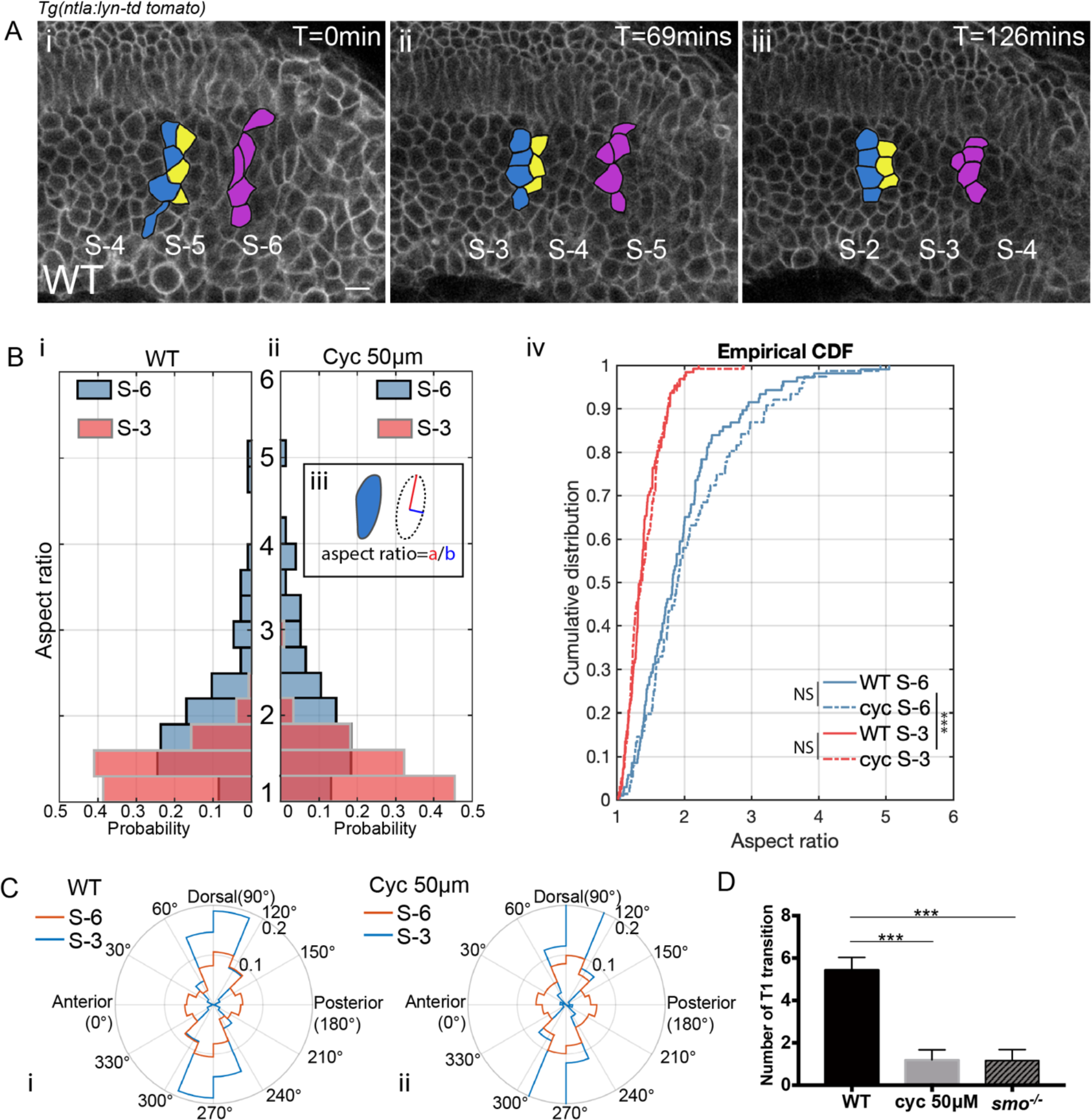
Adaxial cell morphogenesis and intercalations in wild type embryos and embryos under inhibition of Shh signalling. (A) Time lapse of adaxial cell morphogenesis in the posterior PSM. Blue, yellow and violet denote grouped adaxial cells at similar DV but different AP positions. (B) Distribution of the aspect ratios of adaxial cells from wild-type embryos (i) (n_Cells, S-6_=106, n_Cells, S-3_=127, n_Embryos_=4) and embryos treated with cyclopamine at the concentration of 50 μM (ii) (n_Cells, S-6_=76, n_Cells, S-3_=121, n_Embryos_=4). (iii) Cells are fitted by an ellipse that has the same second-moments. The aspect ratio denotes the ratio of the major axis to the minor axis. Blue and red denote cells at stage of S-6 and S-3, respectively. (iv) Cumulative distribution of the aspect ratio of adaxial cells at stage of S-6 and S-3. ***p < 0.001, NS P > 0.05, Kolmogorov-Smirnov test. (C) Distribution of the orientations of the adaxial cell cells from wild-type embryos (i) (n_Cells, S-6_=106, n_Cells, S-3_=127, n_Embryos_=4) and embryos treated with Cyclopamine at 50 μM (ii) (n_Cells, S-6_=76, n_Cells, S-3_=121, n_Embryos_=4). The orientations are obtained according of the direction of the long axis of ellipses fitted from the cells. Blue and red lines denote cells at stage of S-6 and S-3, respectively. (D) The average number of T1 transitions per somite-equivalent region from stage S-5 to S-1 in wild-type embryos (n_Somite_=15, n_Embryos_=5), embryos under Cyclopamine treatment at the concentration of 50 μM (n_Somite_=15, n_Embryos_=5) and *smo^-/-^* mutants (n_Somite_=15, n_Embryos_=5). Images are taken at somites 18-20. Scale bars 20 μm. ***p < 0.001, Student’s t test.

**Supplementary Figure 5.**
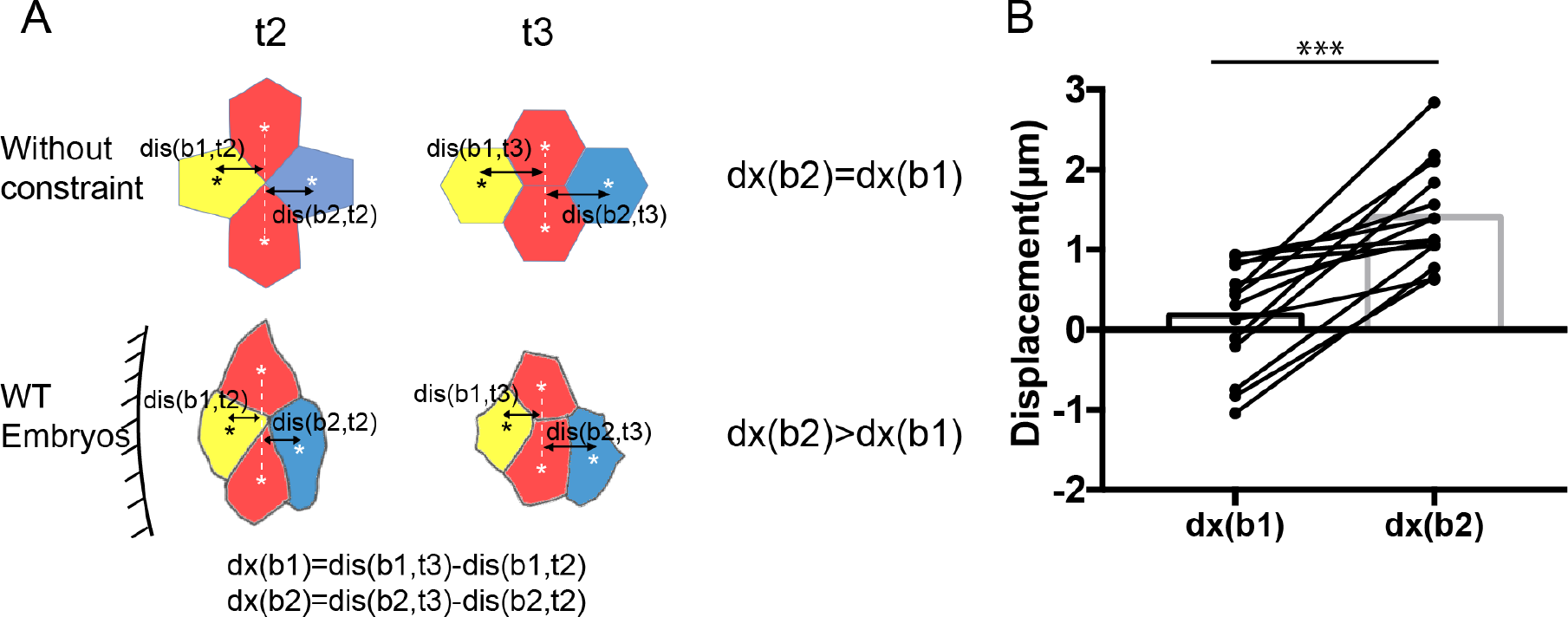
Asymmetric cell extensions under directional cell intercalations. (A) Diagram of idealized T1 transition without external constrain (top panel) and an example of adaxial cell intercalation from wild-type embryos (bottom panel). The relative extensions of cell b1 and cell b2 are measured according to the distance of their centroids to the connecting line (white dashed lines) between the cell a1 and a2 at time point t2 (formation of rosette) and t3 (disjoin of cell b1 and b2). White and black asterisks denote the centroids of cells. (B) Comparisons of the extension of cell b1 towards the anterior direction and the extension of cell b2 towards the posterior direction (n_Intercalations_=14, n_Embryos_=3). ***p < 0.001, Student’s t test.

**Supplementary Figure 6.**
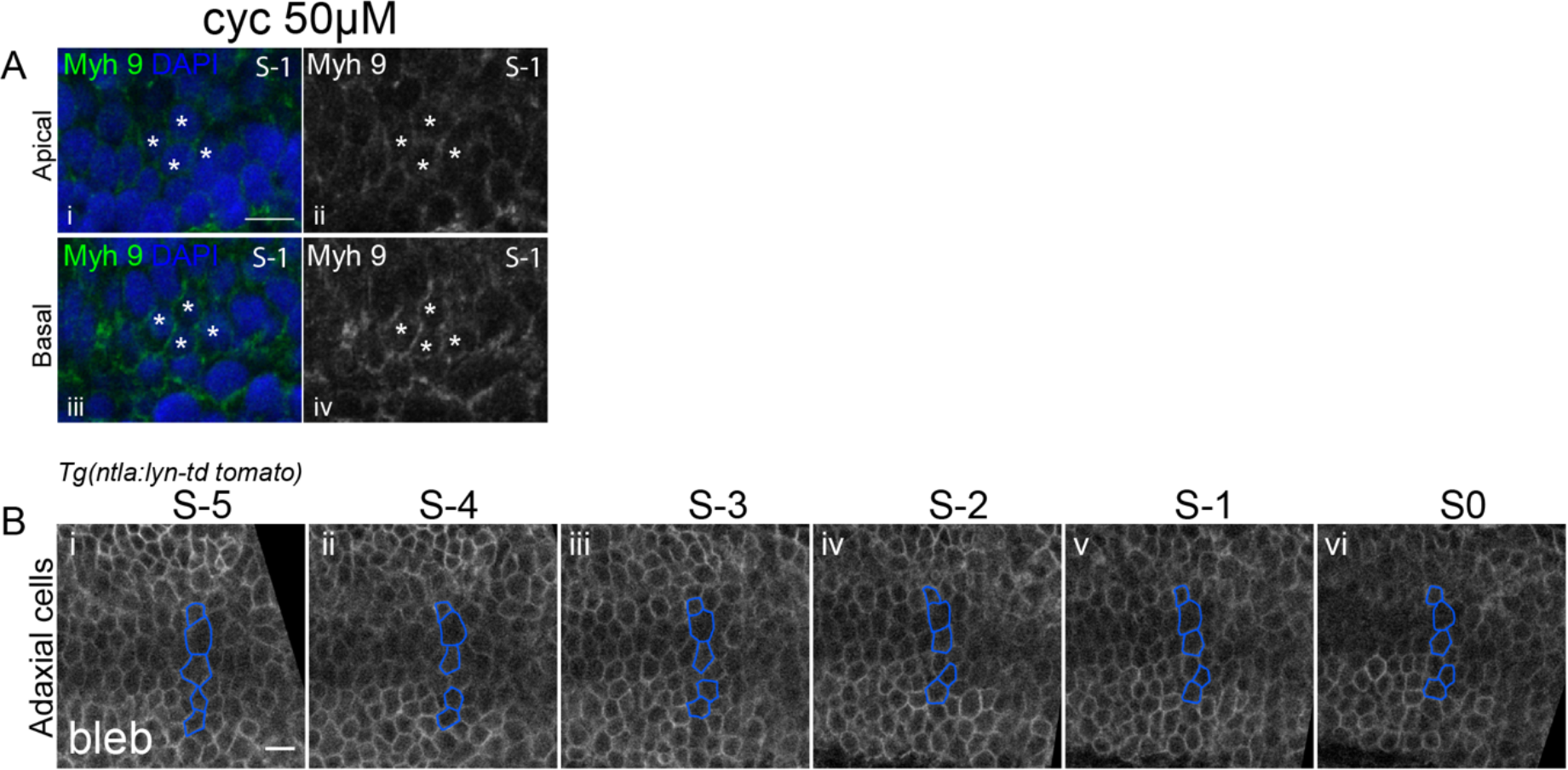
Distribution of Myh9 and effects of blebbistatin treatment. (A) In a subset of adaxial cells from embryos under cyclopamine treatment at 50 μM, stronger intensity of Myh9 is apparent at the basal (iii-iv) instead of apical surface (i-ii). Adaxial cells are labelled with asterisks. White asterisks label adaxial cells at stage S-1. (B) Cell tracking of adaxial cells in embryos treated with blebbistatin at the concentration of 100 μM. Blue color denotes adaxial cells that are localized at similar AP positions at stage S-5 (i). Images are taken at 18-somite stage.

**Movie 1** Cell tracking of adaxial cells located near the anterior somite boundary from stage S-5 to S0. Cell membrane is labelled with Lyn-td tomato. Corresponds to Figure 1E.

**Movie 2** Z-stack of PSM at positions of S0 to S-1 from lateral to medial planes in a wild type embryo. Actin is labelled with Phalloidin 488. Dorsal, DV midline and ventral adaxial cells are labelled with violet, orange and green colours respectively. Red circle labels the apical-to-basal neighbor exchanges. Corresponds to Figure 4D.

**Movie 3** Z-stack of PSM at positions of S0 to S-1 from lateral to medial planes in a *smo*^-/-^ mutant. Actin is labelled with Phalloidin 488. Dorsal, DV midline and ventral adaxial cells are labelled with violet, orange and green colours respectively. Corresponds to Figure 4F.

