## Supplemental Data 1 for "Shh induces symmetry breaking in the presomitic mesoderm by inducing tissue shear and orientated cell rearrangements"

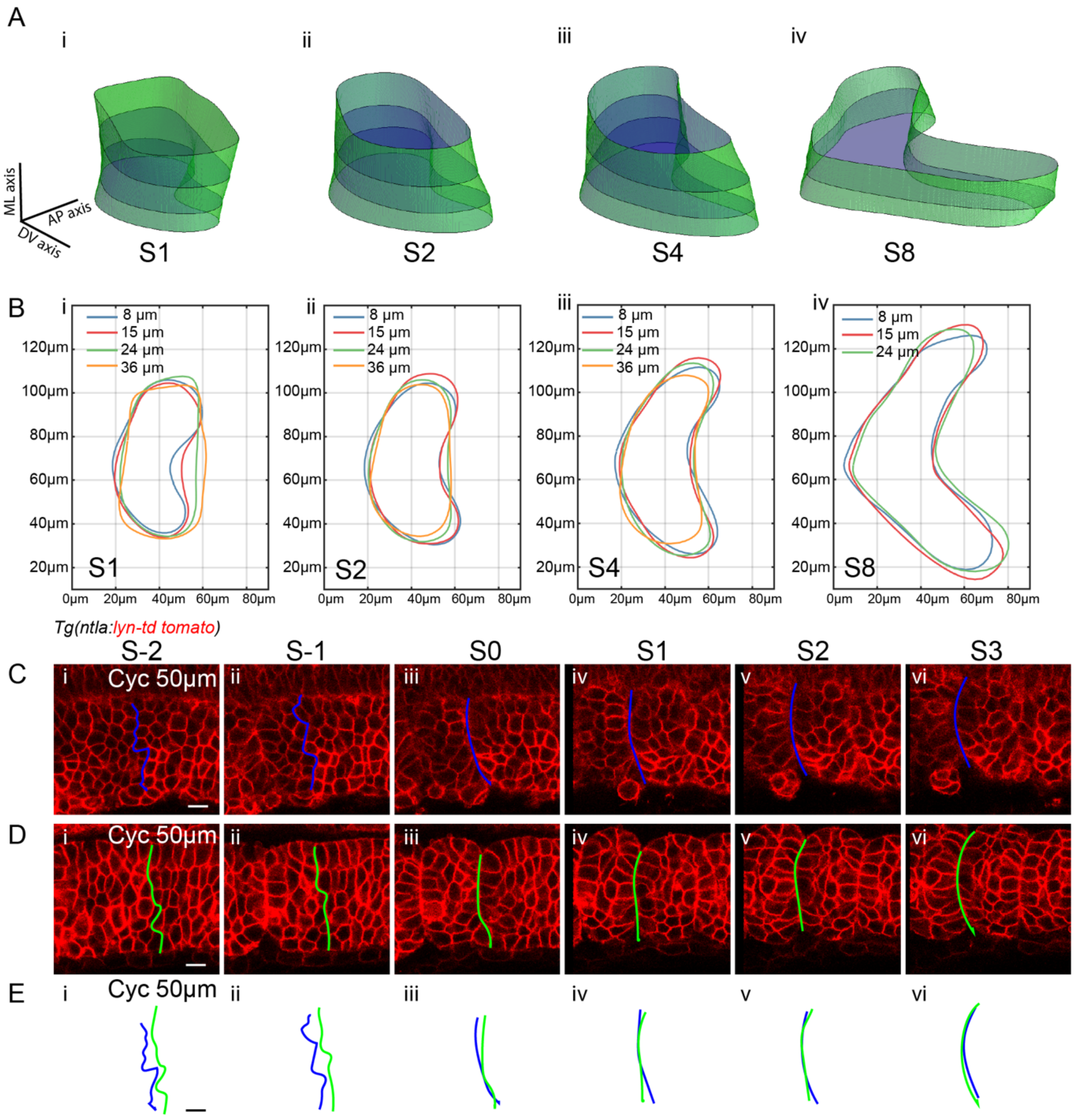


**Supplementary Figure 1. Somite morphogenesis and cell rearrangements under Shh inhibition.**

(A) 3D somite shapes reconstructed from the contours of somites at parasagittal planes 36, 24, 15 and 8 μm from the notochord. (B) Contours of somites at stage S1, S2, S4 and S8. Colors denote the contours at different ML positions. The contours of somites are taken at somite 18. (C-E) Somite boundaries at the adaxial cells and lateral somitic cells from stage S-2 to S3 under 50 μM cyclopamine treatment: (C) adaxial cells; (D) lateral somitic cells; and (E) overlay of the two different boundaries. Images are taken at somites 18-20. Scale bars 10 μm.

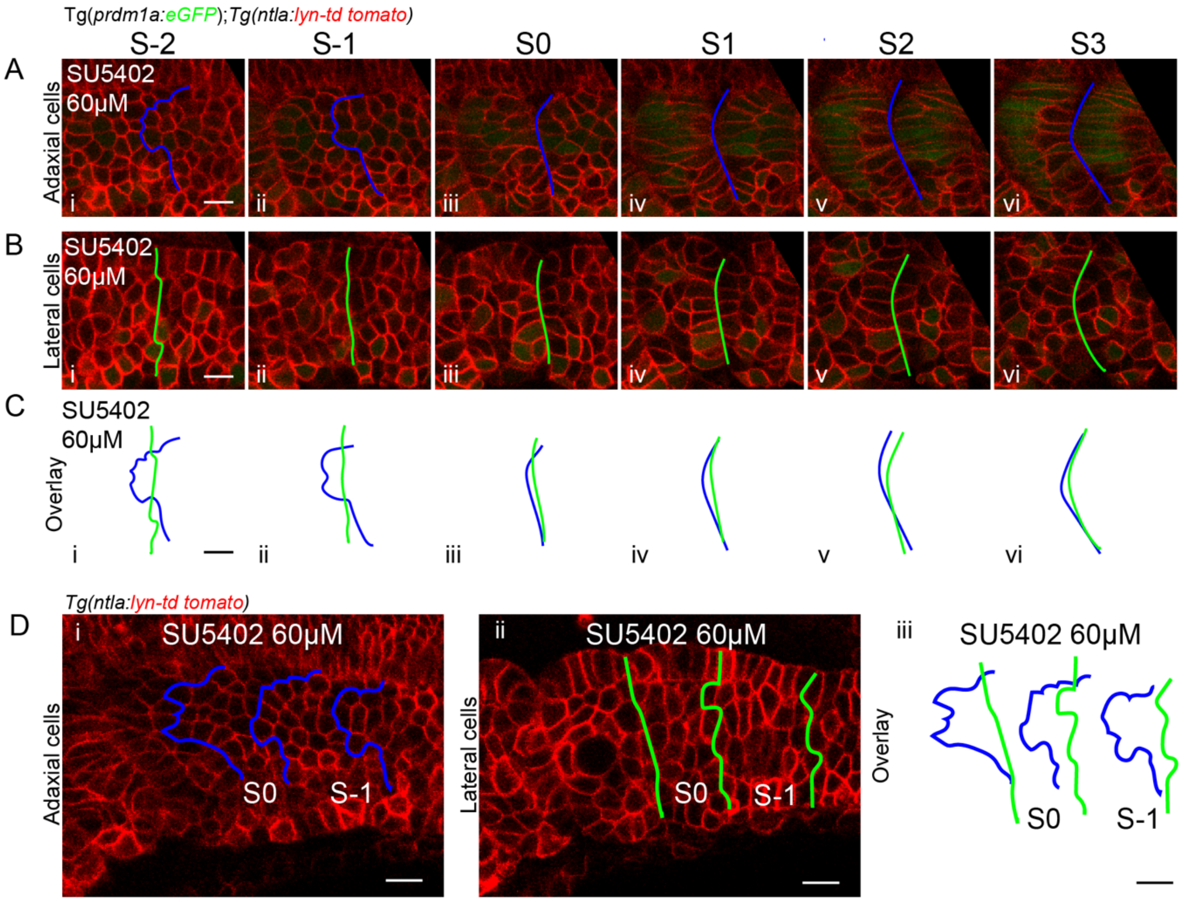


**Supplementary Figure 2. The distinct somite borders at adaxial and lateral somitic cells appear under FGF inhibition**

(A-C) Somite boundaries at the adaxial cells and lateral somitic cells from stage S-2 to S3 under 60 μM SU5402 treatment: (A) adaxial cells; (B) lateral somitic cells; and (C) overlay of the two different boundaries. (D) Relatively small (S0) and large (S-1) somite were induced under 60 μM SU5402 treatment, whilst the distinct somite borders at adaxial and lateral somitic cells occur. Images are taken at somites 18-20. Scale bars 10 μm.

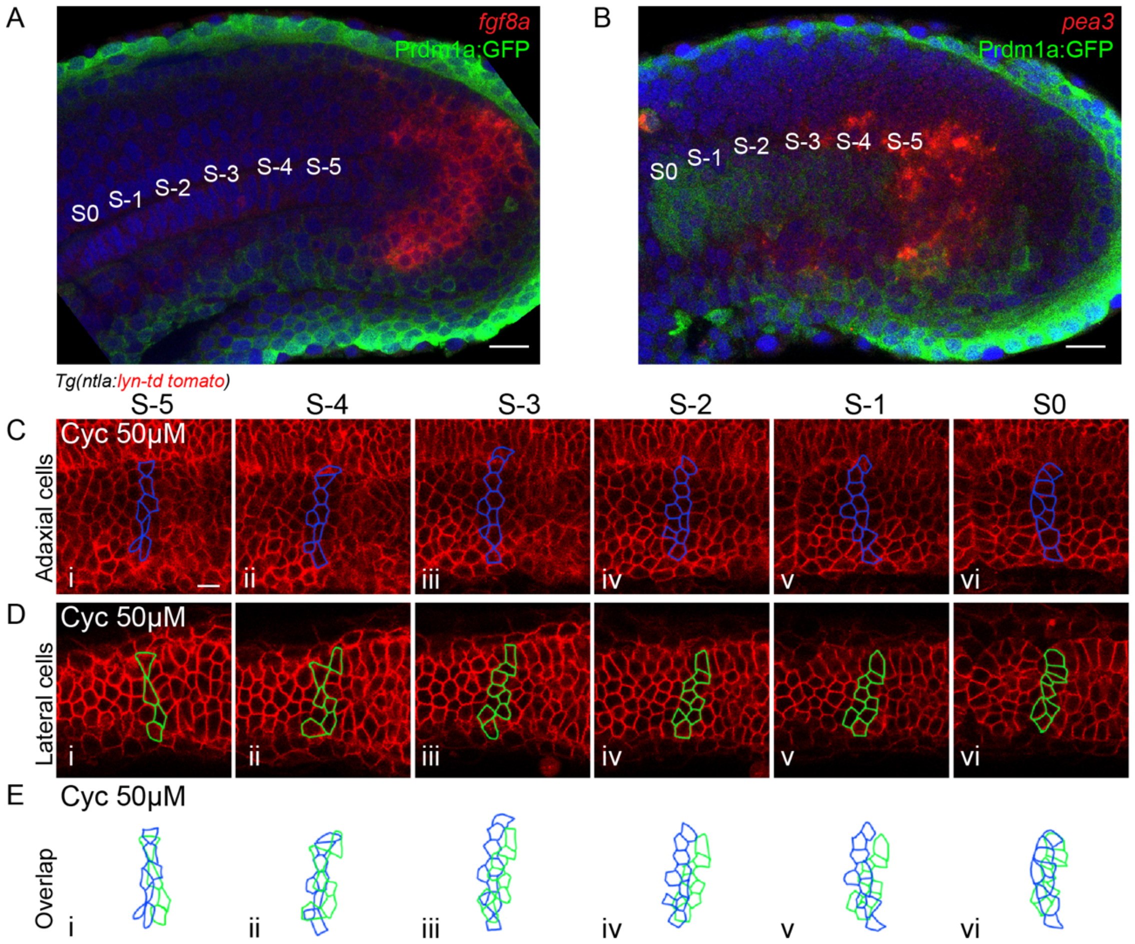


**Supplementary Figure 3. Fluorescent *in situ* of *fgf8a* and *pea3* and cell tracking in the PSM under Shh inhibition**

(A-B) Expression of *fgf8a* (A) and a downstream factor of FGF signalling pathway, *pea3* (B), within the tailbud and posterior PSM. Adaxial cells are labelled with Prdm1:GFP. Images are taken from the parasagittal optical planes at the same ML planes of the notochord and adaxial cells respectively. (C-E) Cell tracking identified the adaxial cells (C) and lateral somitic cells (D) located at the anterior somite border at somite stage S0 (C(vi) and D(vi)) in embryos under 50 μM cyclopamine treatment. Cells are traced back to stages as early as somite stage S-5 (C(i) and D(i)). (E) Overlay of the boundary adaxial cells and boundary lateral somitic cells from (C-D). Images are taken at somites 18-20. Scale bars 10 μm.

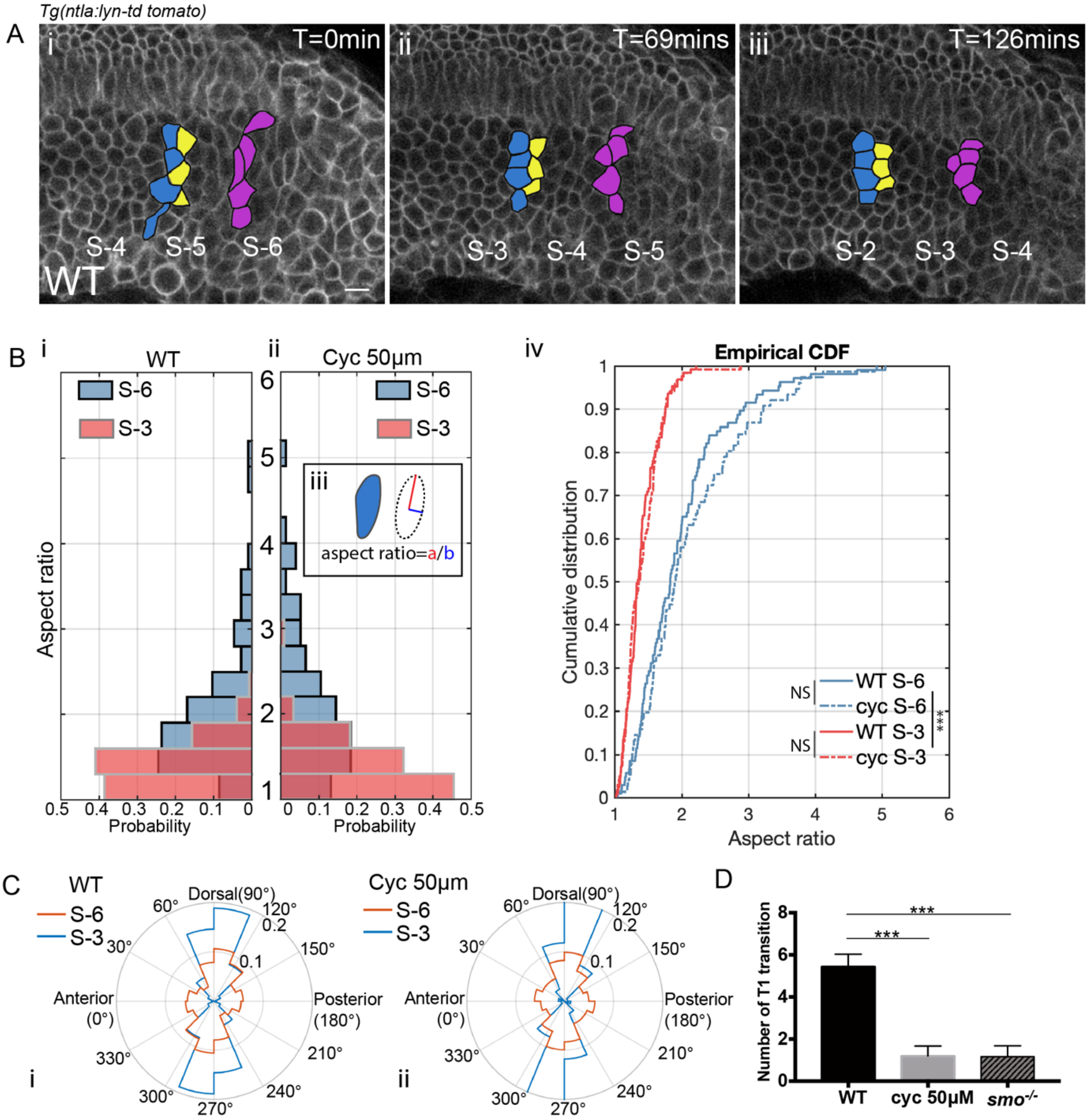


**Supplementary Figure 4. Adaxial cell morphogenesis and intercalations in wild type embryos and embryos under inhibition of Shh signalling.**

(A) Time lapse of adaxial cell morphogenesis in the posterior PSM. Blue, yellow and violet denote grouped adaxial cells at similar DV but different AP positions. (B) Distribution of the aspect ratios of adaxial cells from wild-type embryos (i) ($n_{Cells,\ S\text{-}6}$=106, $n_{Cells,\ S\text{-}3}$=127, $n_{Embryos}$=4) and embryos treated with cyclopamine at the concentration of 50 μM (ii) ($n_{Cells,\ S\text{-}6}$=76, $n_{Cells,\ S\text{-}3}$=121, $n_{Embryos}$=4). (iii) Cells are fitted by an ellipse that has the same second-moments. The aspect ratio denotes the ratio of the major axis to the minor axis. Blue and red denote cells at stage of S-6 and S-3, respectively. (iv) Cumulative distribution of the aspect ratio of adaxial cells at stage of S-6 and S-3. ***$p < 0.001$, NS $P > 0.05$, Kolmogorov-Smirnov test. (C) Distribution of the orientations of the adaxial cell cells from wild-type embryos (i) ($n_{Cells,\ S\text{-}6}$=106, $n_{Cells,\ S\text{-}3}$=127, $n_{Embryos}$=4) and embryos treated with Cyclopamine at 50 μM (ii) ($n_{Cells,\ S\text{-}6}$=76, $n_{Cells,\ S\text{-}3}$=121, $n_{Embryos}$=4). The orientations are obtained according of the direction of the long axis of ellipses fitted from the cells. Blue and red lines denote cells at stage of S-6 and S-3, respectively. (D) The average number of T1 transitions per somite-equivalent region from stage S-5 to S-1 in wild-type embryos ($n_{Somite}$=15, $n_{Embryos}$=5), embryos under Cyclopamine treatment at the concentration of 50 μM ($n_{Somite}$=15, $n_{Embryos}$=5) and *smo*$^{-/-}$ mutants ($n_{Somite}$=15, $n_{Embryos}$=5). Images are taken at somites 18-20. Scale bars 20 μm. ***$p < 0.001$, Student's t test.

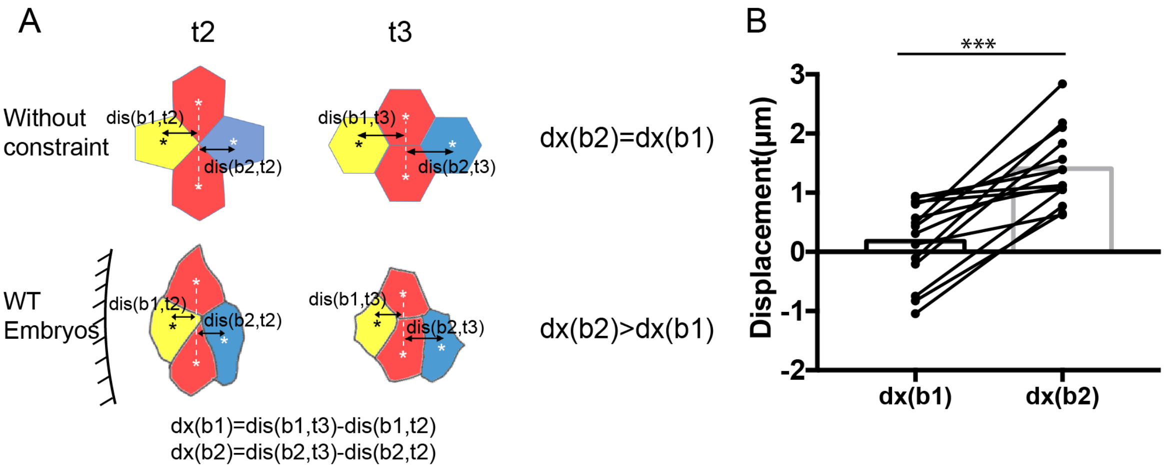


**Supplementary Figure 5. Asymmetric cell extensions under directional cell intercalations.**

(A) Diagram of idealized T1 transition without external constrain (top panel) and an example of adaxial cell intercalation from wild-type embryos (bottom panel). The relative extensions of cell b1 and cell b2 are measured according to the distance of their centroids to the connecting line (white dashed lines) between the cell a1 and a2 at time point t2 (formation of rosette) and t3 (disjoin of cell b1 and b2). White and black asterisks denote the centroids of cells. (B) Comparisons of the extension of cell b1 towards the anterior direction and the extension of cell b2 towards the posterior direction ($n_{Intercalations}$=14, $n_{Embryos}$=3). ***$p < 0.001$, Student's t test.

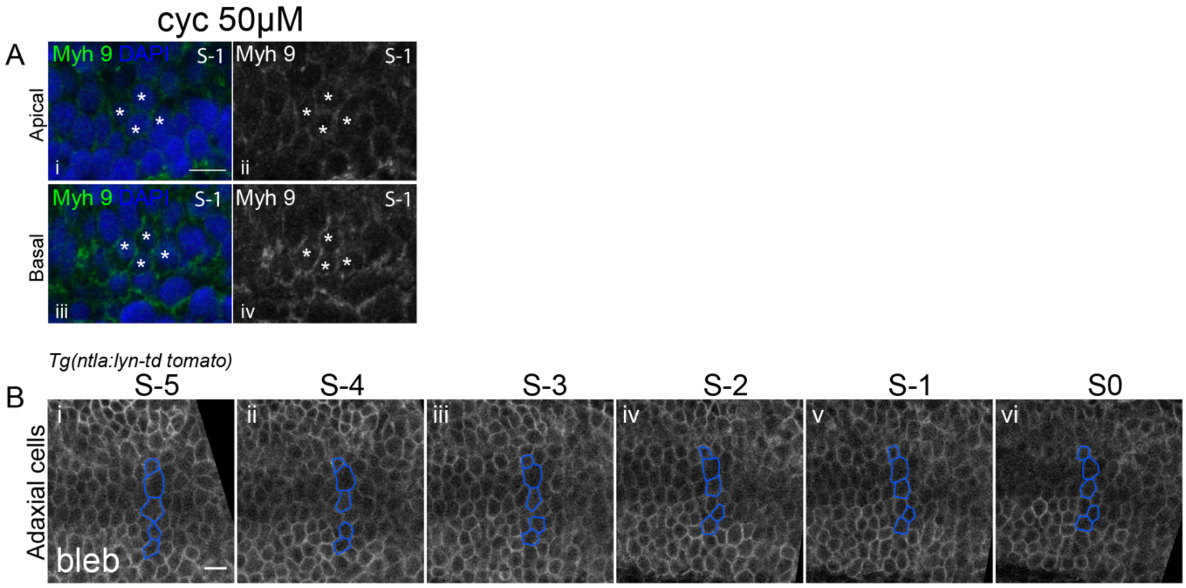


**Supplementary Figure 6. Distribution of Myh9 and effects of blebbistatin treatment**

(A) In a subset of adaxial cells from embryos under cyclopamine treatment at 50 μM, stronger intensity of Myh9 is apparent at the basal (iii-iv) instead of apical surface (i-ii). Adaxial cells are labelled with asterisks. White asterisks label adaxial cells at stage S-1. (B) Cell tracking of adaxial cells in embryos treated with blebbistatin at the concentration of 100 μM. Blue color denotes adaxial cells that are localized at similar AP positions at stage S-5 (i). Images are taken at 18-somite stage.

**Movie 1**

Cell tracking of adaxial cells located near the anterior somite boundary from stage S-5 to S0. Cell membrane is labelled with Lyn-td tomato. Corresponds to Figure 1E.

**Movie 2**

Z-stack of PSM at positions of S0 to S-1 from lateral to medial planes in a wild type embryo. Actin is labelled with Phalloidin 488. Dorsal, DV midline and ventral adaxial cells are labelled with violet, orange and green colours respectively. Red circle labels the apical-to-basal neighbor exchanges. Corresponds to Figure 4D.

**Movie 3**

Z-stack of PSM at positions of S0 to S-1 from lateral to medial planes in a $smo^{-/-}$ mutant. Actin is labelled with Phalloidin 488. Dorsal, DV midline and ventral adaxial cells are labelled with violet, orange and green colours respectively. Corresponds to Figure 4F.
